# Somatic differentiation evolves rapidly and repeatedly through the modification of developmental plasticity

**DOI:** 10.64898/2026.09.22.753194

**Authors:** Dinah R. Davison, Ryan Ruboyianes, Yanghuan Yu, Richard E. Michod, Bradley J.S.C. Olson

## Abstract

Plasticity has been proposed to facilitate the evolution of novelty, but it’s unclear if it can drive a transition to a novel kind of individual. We report the experimental evolution of obligate somatic differentiation in *Eudorina,* a predominantly undifferentiated species that develops plastic somatic cells in response to cold stress. *Eudorina* is a species of volvocine algae, a clade in which cellular differentiation has evolved repeatedly. We exposed *Eudorina* from 2 lab lines and 46 course-based undergraduate research experience lines to repeated cold shock. Somatic differentiation evolved rapidly, with 21 of 48 lines (44%) exhibiting a larger proportion of differentiated colonies in baseline conditions. We also identified a lineage with obligate somatic differentiation; this is the first published example of the experimental evolution of obligate somatic differentiation. 11 lines, including the obligately differentiated one, also exhibited changes in the proportion of somatic cells per differentiated colony. The proportion of somatic cells in these lines converged with that of *Pleodorina,* a polyphyletic soma-differentiated volvocine algae genus. Genome sequencing revealed that 93% of genes mutated in the obligately differentiated line and not in the control had at least one homolog with soma-specific expression in obligately differentiated *Volvox carteri* or *Astrephomene gubernaculifera* and/or cold-induced expression in unicellular *Chlamydomonas reinhardtii.* Moreover, 72% of genes mutated in the obligately differentiated line and not in the control had homologs with both soma-specific expression and cold-induced expression. These genes all have regulatory mutations and lie at the intersection of cold shock and somatic differentiation regulatory modules. Our results provide the first empirical demonstration that the modification of plastic responses to the environment can drive the evolution of somatic differentiation, a key step in the evolution of a new kind of evolutionary individual.

## Introduction

Cellular differentiation is a key step in the evolution of multicellular individuals, a novel evolutionary individual distinct from unicellular individuals. Differentiated multicellularity has evolved repeatedly across the tree of life, including multiple times within green algae alone (Umen and Herron 2021). The repeated evolution of a trait within a clade can result from parallel de novo origins or from changes in the regulation of an ancestrally plastic trait (West-Eberhard 2003). Even though the modification of plastic responses is hypothesized to be an important pathway for the emergence of novelty, relatively few examples exist, and it remains unclear whether it can shape the evolutionary transition to differentiated multicellularity.

We previously proposed that plasticity may have shaped the evolution of multicellular individuality in the volvocine green algae, a clade of freshwater Chlorophyte green algae in which cellular differentiation has evolved repeatedly (Davison et al. 2025; Davison and Michod 2021). The volvocine green algae clade spans a complexity gradient from unicellular genera (e.g., *Chlamydomonas*) to obligate germ-soma-differentiated genera (e.g., *Volvox*), including facultatively differentiated *Eudorina* and obligately soma-differentiated *Pleodorina* and *Astrephomene* (Figure 1A-1F). Somatic cells are small, living cells that do not reproduce and have functional flagella; they have evolved repeatedly in the volvocine algae (Grochau-Wright et al. 2017; Lindsey et al. 2024).

**Figure 1.**
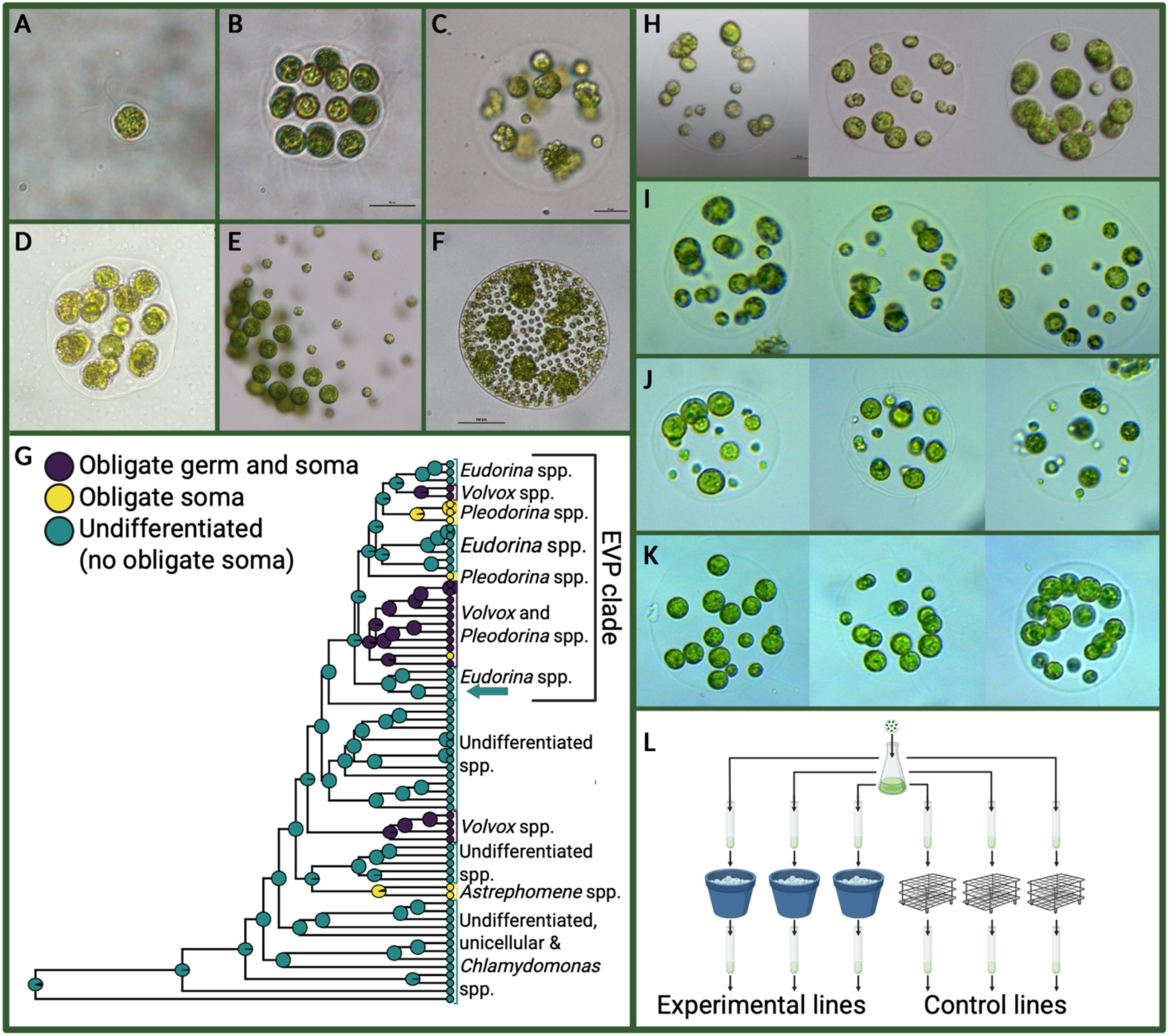
Obligate somatic differentiation has evolved repeatedly in the volvocine algae, including in experimentally evolved *Eudorina.* A-F: Representative images of extant volvocine algae species, modified from Herron (2016) and Davison et al. (2025) with permission. A. *Chlamydomonas reinhardtii.* B. *Eudorina elegans.* C. *E. elegans* following cold shock. Both plastic somatic cells (small cells) and dividing reproductive cells are present. D. *Astrephomene gubernaculifera.* E. *Pleodorina californica.* F. *Volvox carteri.* G. A volvocine algae phylogeny, modified from Lindsey et al. (2024). Species in teal are characterized as undifferentiated (lacking obligate soma, though they may have plastic soma); species in yellow or purple have obligate somatic cells. The arrow shows the phylogenetic position of *Eudorina* sp. NIES 3984. H-K: Three representative colonies per line from four experimentally evolved lines; the small cells are somatic. H. Experimentally evolved *Eudorina* line 11A5. I. Experimentally evolved *Eudorina* from Sec33_Gatos. J. Experimentally evolved *Eudorina* from Sec27_Team3. K. Experimentally evolved *Eudorina* from Sec27_Team5. L. An overview of the cold shock procedure. A flask was created from a single colony, and tubes were set up from that clonal culture. Treatment lines were cold-shocked and then transferred, while control lines were transferred. Created in https://BioRender.com.

Convergent evolution is common in the volvocine algae, and genus-typical morphology has evolved repeatedly. Apart from *Astrephomene,* all the volvocine algae genera listed above are most likely polyphyletic (Lindsey et al 2024). Moreover, all three of the genera in the Eudorina clade (*Eudorina, Pleodorina,* and *Volvox*) are polyphyletic, and obligate somatic cells are hypothesized to have been gained at least three times and lost at least once within this clade (Grochau-Wright et al. 2017; Lindsey et al. 2024) (Figure 1G). The repeated evolution of somatic cells in the Eudorina clade may have been shaped by ancestral plasticity in somatic cell development that was repeatedly modified, resulting in the multiple instances of fixation and loss observed in this clade.

Consistent with this possibility, the delineation between *Pleodorina* and *Eudorina* has historically been contentious (Goldstein 1964; Nozaki et al. 1989), and changes in plastic responses to the environment may have played a role in the repeated evolution of differentiated *Pleodorina* and undifferentiated *Eudorina* morphologies. Grove (1915) found the differentiated species *Pleodorina (Eudorina) illinoisensis* in a puddle covered by ice. After the temperatures warmed and the ice melted, he also observed undifferentiated *Eudorina* and proposed that *Pleodorina* morphology may occur as a physiological form of *Eudorina.* We previously found that *Eudorina elegans* develops plastic somatic cells following two hours of cold shock (Davison et al. 2025), demonstrating that somatic cells can develop in response to environmental conditions. The existence of facultative somatic cells was also documented in other *Eudorina* species in the 1900s (Pascher 1927; W. B. Crow 1925; Hartmann 1921; Iyengar 1933; Smith 1931; W. Bernard Crow 1918). If transitions between predominantly undifferentiated and obligately differentiated morphologies occur through the modification of plastic differentiation, we expect that obligate differentiation should evolve readily from a facultatively differentiated ancestor.

We used experimental evolution to test the hypothesis that cellular differentiation can evolve through changes to ancestral plastic responses. We repeatedly exposed *Eudorina sp.* NIES 3984 to cold shock and characterized the changes in somatic differentiation in 2 lab lines and 46 lines from an introductory biology course-based undergraduate research experience (CURE). We found that the proportion of differentiated colonies significantly increased in 21 of these lines, with 10 lines also evolving increases in the proportion of somatic cells per differentiated colony. Moreover, we subcloned one line and identified a lineage with obligate differentiation (11A5). We sequenced 11A5 and a control and found that somatic differentiation was associated with predominantly regulatory mutations in genes whose homologs are involved in both the cold shock response and somatic differentiation in other species. This is the first study to report the experimental evolution of obligate somatic differentiation.

## Methods

### Experimental evolution: laboratory lines

We used *Eudorina* sp. NIES 3984 (hereafter “*Eudorina”*), a species whose genome has been sequenced and that has been characterized as undifferentiated (Hamaji et al. 2018, 2018). *Eudorina* was kindly provided by Dr. Nozaki, and all cultures were grown at 25⁰C in a 16:8 light:dark cycle. We began by inoculating a flask with a single expanded and undifferentiated colony to control for genetic variation. We let the flask grow for four weeks and then repeatedly withdrew 1.5 mL from the flask and placed it in 20 tubes containing 15mL of Standard Volvox Media (SVM). 10 of the tubes were designated as control lines, and 10 were designated as experimental lines.

After 6 days of growth, lines were transferred to separate culture tubes. 10mL of culture was transferred from each experimentally evolving line to Falcon tubes. The tubes were then placed in an enclosed ice-water bath at 1⁰C for two hours. After two hours of cold shock, the contents of each cold-shocked Falcon tube were transferred to fresh glass tubes with 10mL of SVM (Figure 1L). Since cold shock can kill colonies and slow down growth (Davison et al. 2025), only 5mL from each control line was then placed in tubes with 15mL of fresh SVM.

This process was repeated every five days, corresponding to 2-3 generations. Since 2 hours of cold shock results in substantial mortality (Davison et al. 2025), repeated cold shock selected for survival in response to transient cold temperatures. After 6 rounds of cold shock, all experimentally evolved lines were dead; we therefore returned to the lines generated after 5 rounds of cold shock and maintained the living lines 11A and 19A in the culture collection, along with their corresponding control lines 1A and 9A. We refer to these five rounds of cold shock as “cold stress season 1.” Past selection affecting plasticity can affect the subsequent response to selection. In concordance with this, we implemented a second season of 7 rounds of repeated cold shock (“cold stress season 2”) 7 months (approximately 52-105 *Eudorina* generations) later.

### Characterization of the response to selection

Following the final cold shock treatment, lines were maintained in culture collection for five months (approximately 37 to 75 generations) before being transferred to 125mL replicate flasks containing 30mL of SVM and grown on a shaker for 5-6 days. 20 uL of culture was repeatedly withdrawn and placed on a grid slide with a coverslip for systematic imaging with a Nikon Eclipse TE inverted microscope. 100 expanded adult colonies were characterized by the number of somatic cells they possessed and their estimated total cell number.

To examine whether heritable variation in the development of somatic cells existed within lines and determine whether somatic differentiation was obligate in some lineages, we isolated 6 single differentiated colonies from experimentally evolved lines 11A and 19A and 4 colonies from control lines 1A and 9A. These colonies were placed in separate wells in a 24-well plate and allowed to reproduce for two weeks (3-7 generations) before being examined with a Nikon Eclipse TE inverted microscope. We retained subclones for which all descendant colonies in the well were differentiated. We identified one subclone from 11A, which we named 11A5, in which all the descendant colonies in the well were differentiated. We further characterized this lineage following the characterization procedures described above.

We used R v4.5 for all statistical analyses and visualized data using ggplot2 in R. We used binomial GLMMs with flask as a random effect and Fisher’s exact test to compare the proportion of colonies with somatic cells between experimentally evolved and control lines and between lines of each treatment. We used beta-binomial GLMMs with flask as a random effect and Wilcoxon rank sum tests to examine the proportion of somatic cells per differentiated colony. Multi-panel figures were created in https://BioRender.com

### Experimental evolution: CURE

To determine whether our results were due to the specifics of our selection protocol, we evolved additional *Eudorina* lines as part of a course-based undergraduate research experience (CURE) in the introductory biology lab ECOL 182L at the University of Arizona. Over the course of half a semester, hundreds of undergraduate students across 9 lab sections sought to address whether cellular differentiation could evolve in response to repeated environmental stress.

Students in each lab section were grouped into teams of 2-4 and were assigned one of three treatment protocols. The treatment protocols differed in the frequency and total number of cold shock treatments. All treatment protocols consisted of multiple instances two hours of cold shock and a weekly transfer to fresh media; all teams had a control line as well as an experimentally evolving line. When students were absent, dropped one of the tubes, or mixed up their treatments, they received replacement control and cold treatment tubes from treatment 1 tubes that were independently cold-shocked and transferred each week by ECOL 182L staff.

In treatment 1, students cold-shocked the algae once a week for 5 weeks, for a total of 5 cold shock treatments. In treatment group 2, students cold-shocked the algae every 3-4 days for 5 weeks, for a total of 10 cold-shock treatments. In treatment group 3, students cold-shocked the algae every 3-4 days for 3 weeks, for a total of 5 cold-shock treatments. The three treatments allowed us to determine if the experimental evolution of somatic differentiation was robust to the particulars of the selective regime. Specifically, we examined whether the regulation of somatic cells could evolve if we varied the number of rounds of selection and the spacing between rounds of selection. We also determined if the regulation of cold shock would evolve following only one season of temperature fluctuations.

At least three generations after the last time the treatment groups experienced cold shock, students imaged 50 colonies from their experimentally evolved line and 50 colonies from their control line. Students were instructed to use 10X objectives and to only image adult colonies with large reproductive cells and expanded extracellular matrices. One week prior to this imaging, students practiced identifying plastic somatic cells and colonies of the correct age by imaging *Eudorina* that had been cold-shocked two days prior. Students saved the images of the experimentally evolved and control colonies. Graduate teaching assistants helped students with microscopy.

### Image analysis

We compiled images generated by the undergraduate students and analyzed them to obtain data on the number of reproductive and somatic cells in each colony. We trained image segmentation models from 53 manually annotated images over 100 training epochs using Cellpose 2.0 (Pachitariu and Stringer 2022), classified cells based on relative cell sizes, and manually checked the assignments of all colonies. We detected colonies in images using a custom-trained Cellpose 2.0 model that combined cell clustering with ECM boundary detection. We used a second custom Cellpose mode to segment individual cells within each detected colony. We classified cells as somatic if their area was less than 50% of the maximum cell size and less than 300 pixels. The cells that did not meet this threshold were scored as reproductive.

For each colony, we scored the total cell number, the number of reproductive and somatic cells, the cell size ratio, and a focus score. Since volvocine algae cell division results in cell numbers that follow powers of two and some cells were obscured by other cells and not included in the scored total cell number, we estimated the likely total cell number by rounding the observed cell count to the expected values of 4, 8, 16, or 32. We then calculated the proportion of somatic cells as somatic cells / likely total cell number. We manually reviewed all colonies and adjusted the classifications and counts when needed. We discarded all colonies that were partially out of frame, out of focus, or present in another image. We assigned each colony a confidence score (1 = high, 2 = medium, 3 = low) based on how confident we were that the counts and classification were accurate. We discarded all low-confidence colonies following a sensitivity analysis. We used the data in the image names to assign each colony to the experimental or control group and to determine which treatment protocol they followed. We examined the control data for each team and excluded 6 teams with a proportion of somatic cells in control lines more than 2 standard deviations above the mean and with significantly higher control differentiation (suggesting they mixed up their treatments); we excluded an additional 2 teams that were missing data. The final dataset contained 1,608 control colonies and 1,602 experimental colonies from 46 undergraduate teams. 21 of these teams collected data on treatment 1, 10 completed treatment 2, and 15 completed treatment 3.

We used R v.4.5 as well as SciPy (Virtanen et al. 2020) and statsmodels (Seabold and Perktold 2010) in Python v3.11 to analyze data. We used Fisher’s exact test to compare the proportion of colonies with somatic cells in control and experimental treatments and Fisher’s exact test with Benjamini-Hochberg correction to compare the control and experimental treatments for each line. We analyzed the proportion of colonies with somatic cells with geepack (Højsgaard et al. 2006) in R using generalized estimating equations with a logit link and an exchangeable working correlation, clustering by line (N = 92). We used Wald tests to determine if the response to selection differed across treatment protocols.

We performed Mann-Whitney U tests to compare the proportion of somatic cells per differentiated colony in control and treatment lines. We used beta-binomial GLMMs to model the proportion of somatic cells (somatic cells /likely total cells), with line as random effect. We used this model to estimate the effects of treatment (control vs cold shock) on the proportion of somatic cells per differentiated colony and to examine the relationship between the proportion of colonies with somatic cells and the proportion of somatic cells per differentiated colony. We used ggplot2 (Wickham 2011) in R and matplotlib (Hunter 2007) in Python to visualize the data. Multi-panel figures were compiled in https://BioRender.com.

### Comparison to Pleodorina

We compared the proportion of somatic cells per differentiated colony to obligately differentiated *Pleodorina.* Data on typical colony sizes and the proportion of somatic cells per colony were obtained from published species descriptions and from the Herron et al. (2014) dataset, which was kindly provided by MD Herron. We conducted comparisons with data on *P. starrii* colony size and the proportion and number of somatic cells for control colonies grown in both mixed and still culture collection conditions. Since the proportion of somatic cells per colony in *P. starrii* varies across colonies of different sizes (Herron et al 2014), we only included 16-celled *P. starrii* and *Eudorina* colonies in our analysis. We quantified the proportion of soma per differentiated colony for each CURE and lab line and compared them to *P. starrii.* We also compared the proportion of somatic cells per 16-celled differentiated colony for each line to that of pooled differentiated CURE (N = 272) or lab (N = 53) controls. We conducted comparisons with Wilcoxon rank-sum tests, corrected for multiple testing with the Benjamini-Hochberg procedure.

### Genome sequencing and analysis

We used Oxford Nanopore whole-genome sequencing to identify the mutations that occurred during the experimental evolution of somatic differentiation in the obligately differentiated line 11A5. We grew 11A5 and the control 1A in 500mL flasks and collected tissue via filtration. We flash froze the tissue in liquid nitrogen and sent them to SeqCoast Genomics for sequencing. We used Oxford Nanopore long-read sequencing to capture structural variant changes, as cold shock is known to induce transposable element movement in *Volvox carteri* (Miller et al. 1993; Ueki and Nishii 2008). DNA extraction and sequencing were conducted by SeqCoast Genomics. DNA extraction for both lines was performed using the Qiagen DNeasy 96 Powersoil Pro QIAcube HT Kit (#47021) with MagMAX bead beating (FisherSci #A2351) Bead beating lasted for 5 minutes on a Vortex Genie 2. Library preparation was carried out using the Oxford Nanopore Technologies Native Barcoding Kit (#SQK-NBD114) and Long Fragment Buffer. Sequencing was performed on the PromethION 2 Solo platform with a FLO-PRO114M Flow Cell (vR10.4.1) and a 400bps translocation speed. Basecalling was conducted with Dorado 0.9.0 using the super-accurate basecalling model with barcode trimming enabled. Reads with mean Q ≥ 10 were retained.

All genomic analyses were performed on an Azure virtual machine in the terminal and using Python v3.9 and v3.11. The output FASTQ files were concatenated into a single file for each sample and aligned to the *Eudorina* sp. NIES 3984 reference genome (Hamaji et al. 2018) using minimap2 v2.30 (H. Li 2018) with the Oxford Nanopore preset. The reference genome for *Eudorina sp.* NIES 3984 was 184.03 Mbp with 3,180 scaffolds, 61.03% GC content, and an N50 of 564 kb. The sequenced genome of the experimentally evolved line 11A5 had a mean coverage of 15.71x and a mapping rate of 42.02%; the sequenced genome of the control 1A had a mean coverage of 21.04x and a mapping rate of 61.11%. Samtools v1.19.2 (H. Li et al. 2009) was used for sorting and indexing. Mosdepth v 0.3.6 (Pedersen and Quinlan 2018) was used to calculate coverage. BAM files were sorted for subsequent analyses.

The reference genome was repeat-masked and annotated using the funannotate v1.8.17 pipeline (Palmer and Stajich 2019) following (Takahashi et al. 2023). We built a custom RepeatModeler library with RepeatModeler2 v2.0.7 (Flynn et al. 2020) with LTRStruct and used it to softmask the reference genome with RepeatMasker v4.2.1 (Tarailo-Graovac and Chen 2009). 38.02% of the genome was repeat content. We trained the funannotate pipeline using RNAseq data from a closely related *Eudorina* species, *E. elegans* NIES719 (SRA accession SRR13719258) (Lindsey et al. 2021). We ran Funannotate predict with the BUSCO seed species *Volvox*, an existing *Volvox* GFF from Augustus associated with (Jiménez-Marín et al. 2023), and Augustus/GeneMark-ET + EVM consensus. Funannotate update was used to refine UTR predictions. We generated gene models for 19,846 genes. Annotations were obtained with funannotate iprscan (InterProScan) and funannotate annotate (using Pfam, EggNog, SignalP, MEROPs, dbCAN). BUSCO validation was run with the OrthoDB v10 dataset chlorophyta_odb10 in protein mode; BUSCO completeness was 93.4%.

Variant calling for SNPs was performed with NanoCaller v3.6.0 (Ahsan et al. 2021) with preset ont. NanoCaller filtered ∼30% of raw SNP calls as low quality. Depth and variant frequency were subsequently used for additional SNP filtering. In each line, we only included SNPs with QUAL ≥ 20, read depth of at least 10, VAF ≥ 0.98 (near fixation), and ones that were biallelic. We then assigned each SNP as either sample-specific or shared, depending on its presence/absence in each sample. We assigned SNPs as sample-specific if they were present at a frequency of ≥ 0.98 in one sample and ≤ 0.2 in the other. We assigned SNPs as shared if they were present at a frequency of ≥ 0.98 in both samples. To exclude SNPs that were likely to result from Oxford Nanopore sequencing artifacts, we filtered out SNPs in low-complexity regions with homopolymer runs of ≥ 6 bp, imperfect homopolymers ≥ 10bp, dinucleotide repeats ≥ 10 bp, trinucleotide repeats ≥ 15 bp. Mosdepth coverage was used to filter out SNPs and indels where DP ≤ 10 in at least one sample.

We annotated SNPs using the funannotate gene models we previously generated. We mapped the genomic locations of each SNP to the funannotate GFF file with bcftools v1.19.2 (Genovese et al. 2024). We defined genic SNPs as those 2000 bp upstream or 500 bp downstream of the CDS and intergenic SNPs as those that were ≥ 2 kb from any gene. For each genic SNP, we recorded which genomic region it was in: the promoter (2 kb upstream of the transcription start site), the 3’UTR, the 5’UTR, the CDS, an intron, or within 500 bp downstream of the stop codon. We classified SNPs in tRNA and ncRNA genes as genic_other. For SNPs in the promoter, we recorded the distance from the transcription start site. For intronic SNPs, we recorded the distance from the nearest coding sequence and recorded those within 8bp as being in the splice region. We also used these gene models to score each CDS SNP as synonymous, missense, or nonsense. Variant calling for indels was also performed with NanoCaller (Ahsan et al. 2021). QUAL ≥ 20 and genotype quality ≥ 10 were used for quality filtering. We assigned each indel to one of three categories: 11A5-specific, 1A-specific, or shared, based on their presence/absence in both samples and following the logic laid out above. Since coverage varied across the genome, we used samtools depth to check coverage depth for each indel that was called as present in one sample and absent in the other. We excluded sample-specific indels from the dataset if the other sample had ≤ 10x coverage at that position. We also filtered out indels in hypervariable regions to avoid including ones that were likely to be sequencing artifacts. We excluded indels in homopolymer runs of ≥ 4 bp, microsatellites ≥ 3 tandem repeats, and other low-complexity sequences as described above. This pipeline also produced MNVs.

We annotated indels to the funannotate gene models following the methods described above for SNPs. We identified the genomic regions of each indel and classified the CDS indels as either frameshift or in-frame. This indel identification pipeline also identified MNVs, which we classified as synonymous, missense, or nonsense based on codon changes.

Following Aydin et al. (2025), we used a combination of five structural variant (SV) callers to identify structural variants in 11A5 and 1A. We used Sniffles2 v2.6.3 (Smolka et al. 2024), cuteSV v2.1.3 (Jiang et al. 2020), Delly v1.7.3 (Rausch et al. 2012), SVIM v2.0.0 (Heller and Vingron 2019), and DeBreak v1.0.2 (Y. Chen et al. 2023). We merged SVs into one unified dataset per sample and identified how many callers supported each SV. We retained a SV as sample-specific if it was supported by at least 4/5 callers in one sample and 0 callers in the other sample. We retained a SV as shared if it was supported by at least 4/5 callers in both samples. We excluded BNDs calls within 1kb of scaffold terminals, since those were likely to be assembly artifacts. We also excluded SVs in repeat dense regions (2 kb windows with ≥ 60% repeat density). We annotated the SVs with the funannotate gene models using the genomic regions described above. We categorized SVs by type: insertion, deletion, duplication, inversion, and breakend/translocation.

We had additional annotation steps to characterize TE insertions, which we recorded as TEs rather than SVs. We identified TE class and family using a combination of five tools: Dfam nhmmer v3.4 (Wheeler et al. 2013) with 300 Viridiplantae curated HMMs, TEsorter v1.5.1 (R.-G. Zhang et al. 2022) with REXdb Viridiplantae, RepeatModeler2 v2.0.7 (Flynn et al. 2020) Blastn with a *Eudorina-*specific library that we constructed, BLASTX vs RepeatPeps with a TE protein database, and Pfam HMMER v3.4 with TE protein domains. We mapped TEs to the funannotate gene models and included the genomic regions described above. We used TE presence/absence across samples to classify TEs as sample-specific or shared.

We created sample-specific and shared mutation datasets of all genomic changes following the methods above. We searched the datasets to identify genes that were present in one sample-specific dataset and absent from the other sample-specific dataset and the shared dataset. We refer to these gene sets as sample-specific genes. We further filtered out genes that encoded tRNAs, genes we were unable to annotate, genes with only synonymous mutations, and genes with only deep intronic SNPs or indels.

### Sample-specific gene functional annotations

We primarily obtained gene annotations by using NCBI BLASTp and examining the putative conserved domains. For sequences for which conserved domains weren’t identified, we obtained annotation information from volvocine homologs found via NCBI and Phytozome (Goodstein et al. 2012) BLASTp. We supplemented our understanding of gene functions by using the gene annotations generated by the funannotate pipeline. Supplementary material S1 contains additional annotation information.

### Comparisons to other volvocine species

Cell-type-specific gene expression was previously published for *Astrephomene gubernaculifera* (hereafter “*Astrephomene*”) (Yamashita et al. 2021) and *Volvox carteri* (hereafter “*Volvox”*) (Matt and Umen 2017). We used these datasets in conjunction with the published *Volvox* and *Astrephomene* genomes and cell-type specific transcriptomes to determine if some of the same genes and pathways involved in somatic cell development in obligately differentiated volvocine algae species were mutated in 11A5. We used BLASTP v2.11.0+ on Phytozome to identify the *Volvox* homologs of each sample-specific mutated gene and cross-referenced the gene IDs of homologs with bitscores > 50 to the list of genes with soma-specific expression generated by (Matt and Umen 2017). We used NCBI BLASTp to identify *Astrephomene* homologs of each sample-specific mutated gene and cross-referenced them against the published dataset of *Astrephomene* cell-type-specific expression.

We examined whether genes mutated in 11A5 and 1A were part of a volvocine cold shock response module. Gene expression following up to one hour of cold shock was previously published for unicellular *Chlamydomonas reinhardtii* (hereafter “*Chlamydomonas*”) (L. Li et al. 2020). We used BLASTP v2.11.0+ on Phytozome to identify all *Chlamydomonas* homologs with bitscores > 50 of each sample-specific mutated gene and cross-referenced the gene IDs to the dataset published by (L. Li et al. 2020). We used Fisher’s exact test to compare the proportion of sample-specific annotated genes whose homologs showed cold-induced and/or somatic-specific expression in 11A5 vs in 1A. We restricted this comparison to genes with homologs that were identified in all three species. Since 11A5 and 1A are single lines, these tests describe the contrast between the experimentally evolved and control genomes.

## Results

### Experimental evolution: lab lines

#### Proportion of colonies with somatic cells

The proportion of colonies with soma increased following repeated cold shock (Figure 2A). Five months (approximately 37-75 generations) after the last application of cold shock, the repeatedly cold-shocked lines contained significantly more differentiated colonies than did controls (binomial GLMM with flask as a random effect, odds ratio = 4.41, 95% confidence interval = 3.14 – 6.18, p < 0.0001). 43.0% (174/405) of experimentally evolved colonies had one or more somatic cells, while only 14.6% (59/404) of controls were differentiated. The two experimentally evolved lines did not have significantly different proportions of colonies with somatic cells (p > 0.05); 44.3% of colonies in line 11A and 41.6% of colonies in line 19A were differentiated. Similarly, the two control lines did not have significantly different proportions of colonies with soma (p > 0.05); 14.9% of colonies in 1A and 14.4% of colonies in 9A were differentiated.

**Figure 2.**
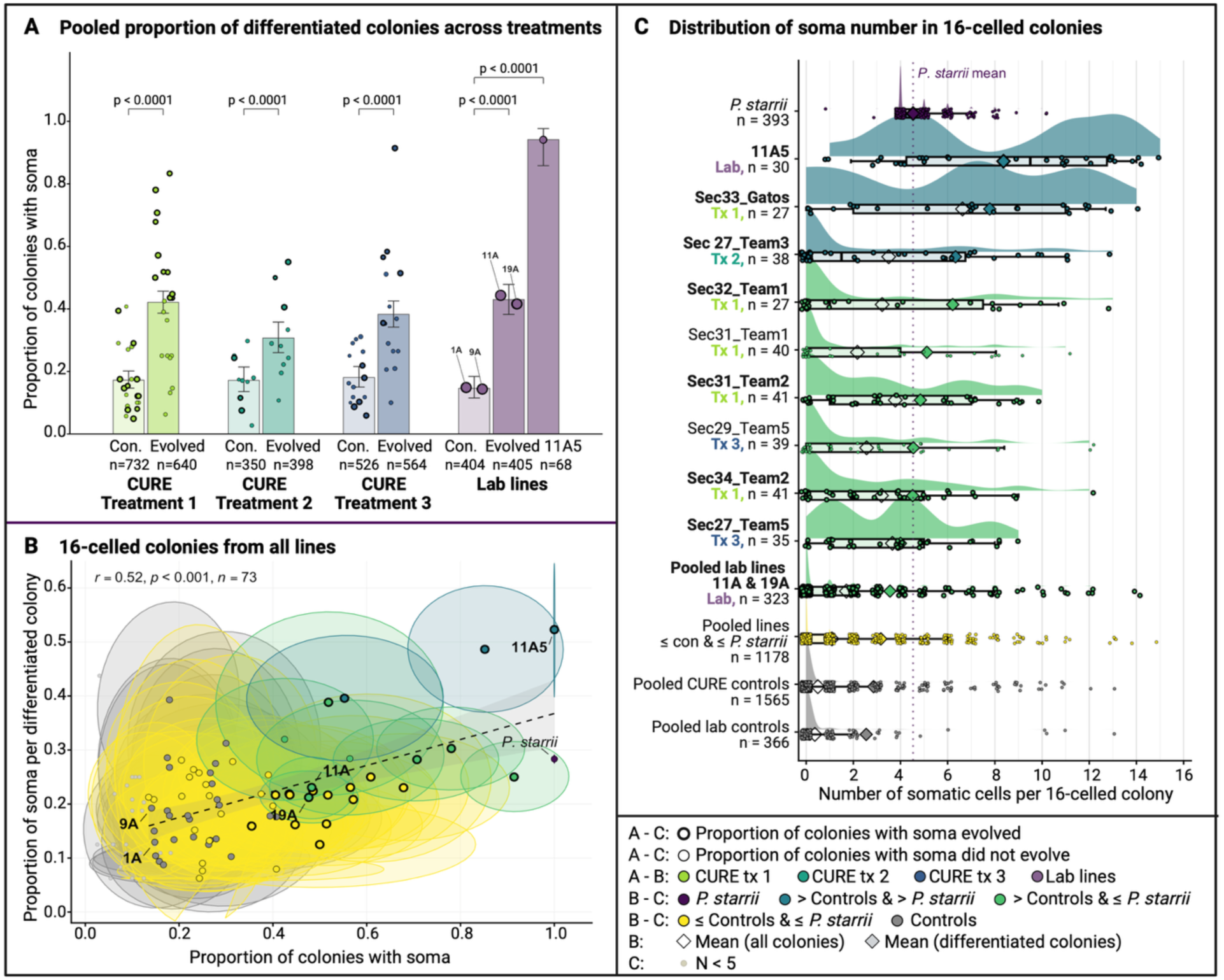
Repeated evolution of somatic differentiation in experimentally evolved Eudorina. **A.** Proportion of colonies with at least one somatic cell in pooled evolved and control lines for each CURE treatment and for the lab lines; obligately differentiated subcloned lineage 11A5 is shown separately. Each bar shows the proportion of colonies with soma; the circles show per-line means. **B.** The proportion of somatic cells per differentiated 16-celled colony plotted against the proportion of colonies with soma. Each point is one line with its 95% confidence interval ellipse; 16-celled *P. starrii* is shown in purple for comparison. The dashed line is the linear fit across *Eudorina* lines with at least 5 differentiated colonies (r = 0.52, p < 0.0001, n = 73 lines). **C.** The distribution of somatic cell number in 16-celled colonies for select lines along with *P. starrii,* pooled controls, pooled lab lines, and pooled CURE lines in which the proportion of soma per colony did not increase. Open diamonds show the mean across all colonies; filled diamonds show the mean across differentiated colonies. The dotted line is the *P. starrii* mean. In panels B and C, color denotes whether the proportion of somatic cells per differentiated colony increased (see key). In all panels, bold labels and points outlined in bold denote lines where the proportion of colonies with soma was significantly greater than paired controls. Created in https://BioRender.com.

#### Proportion of somatic cells per differentiated colony

The regulation of somatic cell development also evolved following repeated cold shock, as the proportion of somatic cells per differentiated colony increased (Figure 2B-2C). Differentiated colonies in the experimentally evolved lines had a higher proportion of somatic cells than did differentiated colonies in the controls (beta-binomial GLMM with flask as a random effect, odds ratio = 1.46, 95% confidence interval = 1.09 – 1.96, p < 0.05). In the experimentally evolved lines, 22.9% of cells in differentiated colonies were somatic, whereas only 15.56 of cells in differentiated control colonies were somatic. The two experimentally evolved lines did not differ (11A = 23.8%, 9A = 18.3%, p > 0.05), nor did the two controls (1A = 12.9%, 9A = 18.3%, p > 0.05). 60% of differentiated 1A colonies and 48.3% of differentiated 9A colonies had only one somatic cell, whereas 34.4% of 11A and 34.5% of 19A colonies had only one somatic cell.

#### Subcloning

To assess the presence of genetic variation in the experimentally evolved lines, single differentiated colonies were transferred to separate wells and allowed to reproduce for two weeks before being examined under a microscope. In 9 of the 10 wells, both differentiated and undifferentiated colonies were present. In a well derived from line 11A, all 25 colonies had somatic cells, and an average of 39.9% of cells were somatic. This line was named 11A5 (Figure 1H) and used in subsequent analyses, including genome sequencing analysis.

To further characterize this line, 11A5 and the corresponding control line 1A were transferred to replicate flasks of SVM 9 months after the last round of cold shock and allowed to grow for one week before imaging. We found that 94.1% (64/68) of 11A5 colonies were differentiated (Figure 2A). All 32 11A5 colonies of 16 or 32 cells were differentiated; the only four colonies scored as undifferentiated were 8-celled. In contrast, 20.9% (9/43) of 1A colonies in corresponding replicate flasks were differentiated; the two lines were significantly different (Fisher’s exact test, odds ratio = 56.6, 95% confidence interval = 15.6 – 273, p < 0.0001). The mean proportion of somatic cells per differentiated colony in 11A5 was 0.41 (median = 0.34, IQR = 0.19 – 0.64). In 1A, the mean proportion of somatic cells per differentiated colony was 0.076 (median = 0.063, IQR = 0.063 – 0.063); 8 of the 9 differentiated colonies contained one somatic cell. The proportion of somatic cells per differentiated colony was significantly higher in 11A5 than in 1A (beta-binomial GLM, odds ratio = 4.25, 95% confidence interval = 1.93 – 9.34, p < 0.001) (Figure 2B – 2C). Additional information is available in supplementary material S1.

### Experimental evolution: CURE

#### Proportion of colonies with somatic cells

To determine the extent to which these results are repeatable and robust to the particulars of our selection protocol, we analyzed the data from the undergraduate CURE lines separately from the lab lines. Repeated cold shock during the CURE increased the proportion of colonies with somatic cells. Across the 46 experimentally evolved lines and paired controls, the proportion of colonies with soma increased from 17.5% (281/1,608) in controls to 38.5% (616/1,602) in experimentally evolved lines, a significant increase of 21.0% (Fisher’s exact test, p < 0.0001). We fit a GEE logistic regression analysis with the 92 lines (from 46 teams) as the clustering unit to account for population structure and found that repeated exposure to cold shock increased the odds of somatic differentiation (odds ratio = 2.98, 95% CI 2.21-4.02, z = 7.15, p < 0.0001), with a within-line correlation of 0.082.

All three treatments had a significantly higher proportions of colonies with soma than their corresponding controls (GEE logistic regression clustering on line, N = 92 lines: T1 odds ratio = 3.49, 95% confidence interval = 2.16 – 5.64, p < 0.001); T2 odds ratio = 2.18, 95% confidence interval = 1.30 – 3.64, p < 0.01; T3 odds ratio = 2.90, 95% confidence interval = 1.74 – 4.84, p < 0.001) (Figure 2A). In treatment 1, 42.1% (315/748) of experimentally evolved colonies and 17.2% (126/732) of control colonies were differentiated. In treatment 2, 30.7% (104/339) of experimentally evolved colonies and 17.1% (60/350) of control colonies were differentiated. In treatment 3, 38.3% (197/515) of experimentally evolved colonies and 18.1% (95/526) of control colonies were differentiated. The magnitude of the response to selection did not differ significantly across treatments when accounting for population structure (condition x treatment interaction, GEE logistic regression clustering on line: Wald χ²₂ = 1.78, p > 0.05).

Individual lines did not all have the same response to selection. The proportion of colonies with soma increased in 78.3% (36/46) of experimentally evolved lines relative to their paired controls; 41.3% (19/46) of these increases were significant after FDR correction (Benjamini-Hochberg, α = 0.05). None of the lines exhibited significant decreases in the proportion of colonies with soma. In treatment 1, 11/21 teams had significant increases in the proportion of differentiated colonies; 3/10 treatment 2 teams showed significant increases, and 5/15 treatment 3 teams showed significant increases in the proportion of differentiated colonies.

#### Proportion of differentiated cells per colony

The proportion of somatic cells per differentiated colony was significantly higher in experimentally evolved lines than in controls (Mann-Whitney U = 70966.5, p < 0.0001), an effect that remained when accounting for population structure (beta-binomial GLMM with line as a random effect, OR = 1.29, 95% confidence interval = 1.07 – 1.56, p < 0.01) (Figure 2B-2C). In experimentally evolved colonies, an average of 24.0% of cells were somatic in differentiated colonies (median = 18.8%, N = 616), while in controls, a mean of 17.9% of cells were somatic in differentiated colonies (median = 12.5%, N = 281). Variation across lines accounted for a small portion of the total variation (ICC = 2.7%), indicating that this difference was not driven by a subset of outlier lines.

The magnitude of this effect depended on whether the proportion of differentiated colonies in the population increased (condition x response-status interaction, likelihood-ratio test χ²_1_ = 5.95, p < 0.05). When we examined only colonies from the 19 lines in which the proportion of colonies with soma increased significantly, we found that the mean percentage of somatic cells per differentiated colony was 26.5% (median = 18.8%, N = 352), compared with 15.8% (median = 12.5%, N = 100) in the corresponding controls, a statistically significant increase (odds ratio = 1.67, confidence interval = 1.26 – 2.22, p < 0.001).

Among the 27 lines in which the proportion of colonies with soma did not change significantly, the mean proportion of somatic cells per differentiated colony was 20.6% (median = 12.5%, N = 264), which was not significantly different from the corresponding controls’ mean of 19.0% (median = 12.5%, N = 181) (odds ratio = 1.06, 95% confidence interval = 0.83 – 1.34, p > 0.05). Experimentally evolved, differentiated colonies from teams where the proportion of colonies with soma evolved had significantly more somatic cells than did colonies from teams in which the proportion of colonies with soma didn’t evolve (odds ratio = 1.33, 95% confidence interval = 1.06 – 1.66, p < 0.05); their corresponding controls did not differ (odds ratio = 0.84, 95% confidence interval = 0.62 – 1.12, p > 0.05).

Across lines, the proportion of colonies with soma predicted the proportion of somatic cells per differentiated colony (beta-binomial GLMM with line as a random effect, OR = 1.11 per 10% increase in differentiation frequency, 95% confidence interval = 1.06 – 1.17, p < 0.0001) (supplementary figure S1). The proportion of colonies with somatic cells explained approximately 37% of the variance in the proportion of soma per differentiated colony among lines. Differentiation frequency was a strong predictor after accounting for experimental treatment (control or cold shock) (likelihood ratio test χ²= 15.5, p < 0.0001), whereas treatment did not explain any additional variation in the proportion of somatic cells per differentiated colony beyond what was explained by differentiation frequency (likelihood ratio test χ²= 0.03, p > 0.05). Supplementary material S1 provides additional information.

### Comparison to Pleodorina

In 11 CURE and lab strains, the proportion of somatic cells per 16-celled differentiated colony increased relative to controls and converged with *Pleodorina* (Figure 2B-2C; supplementary figures S5-S9). The mean proportion of somatic cells of 16-celled *P. starrii* was 0.28 (N = 393). Three lines, including 11A5, evolved a significantly higher proportion of somatic cells than *P. starrii* (11A5: mean soma = 0.52, N = 30, p-adjusted < 0.0001, Figure 1H; Sec33_Gatos: mean soma = 0.49, N = 23, p-adjusted < 0.0001, Figure 1I; Sec 27_Team3: mean soma = 0.40, N = 21, p-adjusted < 0.01, Figure 1J). The proportion of soma per 16-celled differentiated colony for these three lines falls above *P. starrii* and is similar to the standard range for *P. californica* (0.38 – 0.50 (Shaw 1894)). These three lines also evolved significant changes in the proportion of 16-celled colonies with soma (11A5: proportion of 16-celled colonies with soma = 1; Sec33_Gatos: proportion of 16-celled colonies with soma = 0.85, Sec27_Team3: proportion of 16-celled colonies with soma = 0.55).

A further 8 lines evolved a significantly higher proportion of somatic cells per 16-celled differentiated colony than controls, though not significantly more than *P. starrii* (Sec32_Team1: mean soma = 0.39, N = 14, p-adjusted < 0.05; Sec31_Team1: mean soma = 0.32, N = 17, p-adjusted < 0.05; Sec31_Team2: mean soma = 0.30, N = 32, p-adjusted < 0.001; Sec29_Team5: mean soma = 0.28, N = 22, p-adjusted < 0.05; Sec34_Team2: mean soma = 0.28, N = 29, p-adjusted < 0.01; Sec27_Team5: mean soma = 0.25, N = 32, p-adjusted < 0.05, Figure 1K; 11A: mean soma = 0.23, N = 84, p-adjusted < 0.05; 19A: mean soma = 0.21, N = 71, p-adjusted < 0.05) (Figure 2B-2C). These lines also exhibit convergence with *Pleodorina* (supplementary table S11). The species-typical proportion of somatic cells for *P. starrii* ranges from 0.25 to 0.375 (Nozaki et al. 2006); 5 lines fall within this range. Sec32_Team1 falls slightly above this and into the range for *P. californica,* and both lab lines fall within the typical range for *P. japonica* (0.195 to 0.266) (Nozaki et al. 1989).

### Genomic changes

#### Genic changes

Mutations that were only present in one line affected 98 genes in 11A5 and 208 genes in 1A. We focused our analyses on genes with mutations in only one sample. 41 genes were mutated in experimentally evolved 11A5 and not in the control 1A, and 57 genes were mutated only in 1A (supplementary table S16). Of the 41 genes mutated only in 11A5, 23 were excluded as described in the methods, resulting in 18 11A5-specific annotated genes. Of the 57 genes mutated only in 1A, 26 were excluded as described in the methods, resulting in 31 1A-specific annotated genes. Of the 18 11A5-specific genes, 5 (5/18, 27.78%) had CDS mutations (1 of which only had synonymous SNPs), 4 had promoter mutations, 5 had UTR mutations, 5 had downstream mutations, and 8 had intronic mutations. 5 genes had mutations in multiple genic regions. 94.44% (17/18) of these genes had at least one mutation in a regulatory region, demonstrating that the transition to obligate differentiation was largely shaped by regulatory changes. However, the proportion of sample-specific genes with at least one mutation in a regulatory region was not significantly higher in 11A5 than in 1A (Fisher’s exact test, odds ratio = 2.519, p > 0.05). In 1A, 87.10% of genes had at least one mutation in a regulatory region. 9 genes (9/31, 29.03%) had mutations in the CDS, 1 of which had only a synonymous SNP. Additional details are available in supplementary material S1, including supplementary tables S12-S16.

Since obligate differentiation evolved in response to repeated cold shock, the annotations were compared to the known cold stress response pathways in *Chlamydomonas* and *Arabidopsis* (summarized in Ermilova (2020)) (Figure 3A). The functional annotations of 11 11A5 genes mapped onto this cold stress response pathway. The remaining 7 annotated genes fell into two additional clusters: a cell cycle regulation cluster (5 genes) and a cell wall remodeling cluster (2 genes). Although Ermilova (2020) does not treat cell cycle regulation as a separate part of the cold shock response, *Chlamydomonas* cells can cease progression through the cell cycle and division in response to cold stress (Ermilova 2020; L. Li et al. 2020); we therefore added cell cycle regulation to Figure 3A. We excluded cell wall remodeling because control 1A carries a mutation in a different peptidase M11 gene, suggesting that mutations in cell wall genes could reflect adaptation to culture collection conditions rather than to cold shock. We found that mutations in sample-specific mutated genes occurred throughout the cold shock response pathway, with the most changes occurring as part of the protective response downstream of transcriptional regulation (Figure 3E).

**Figure 3.**
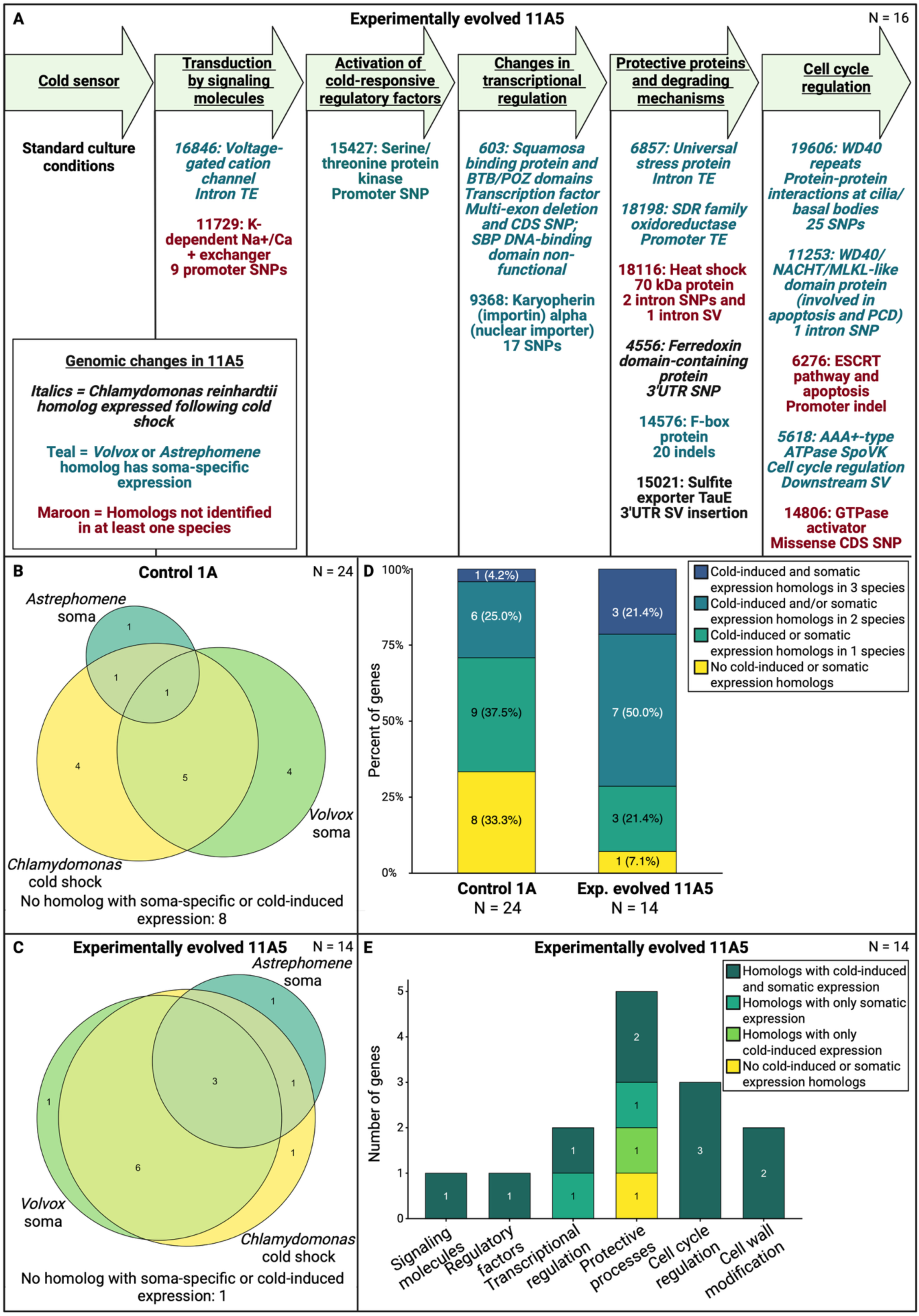
Genes with mutations only in experimentally evolved 11A5 fall disproportionately on genes whose homologs have both cold-induced and soma-specific expression in other species. A. 16 of the 18 annotated genes with mutations only in 11A5 mapped to the stages of the cold stress response pathway or to a cell cycle regulation module. The gene name prefix EUDS and leading zeros are removed. Color and italics correspond to the expression of homologs (see key). B-C. The overlap among genes whose homologs show cold-induced expression in *Chlamydomonas,* soma-specific expression in *Volvox,* and soma-specific expression in *Astrephomene* for 1A (B) and 11A5 (C). Only genes with homologs identified in all three species are included. Genes whose homologs lacked soma-specific or cold-induced expression are counted below each diagram. D. The percentage of genes in each line by the number of species in which a homolog is cold-induced and/or soma-specific. E. The 14 11A5 genes with identified *Chlamydomonas, Volvox,* and *Astrephomene* homologs mapped to the cold stress pathway or to the cell cycle regulation and cell wall modification modules. Colors correspond to homolog expression (see key). Created in https://BioRender.com.

One of the two genes associated with changes in transcriptional regulation is EUDS_000603, which encodes a transcription factor with a multi-exon deletion that spans the DNA-binding domain. Gene EUDS_000603 has a Squamosa Promoter-Binding Protein-Like (SPB) domain and a BTB/POZ domain. Squamosa Promoter-Binding Protein-Like proteins are plant-specific transcription factors that act as developmental regulators, mediating a range of developmental processes, including the transition between vegetative and reproductive growth and responses to environmental stress (X. Chen et al. 2010; Birkenbihl et al. 2005). BTB/POZ domains can mediate dimerization and transcriptional repression (Stogios et al. 2005). The *Chlamydomonas* homolog of EUDS_000603 is differentially downregulated in response to 1 hour of cold shock (L. Li et al. 2020). Therefore, a developmental regulator that may be downregulated during the cold shock response was lost. Additional information about EUDS_000603 is available in supplementary material S1 and supplementary figure S10.

### Comparisons to other volvocine species

We examined sample-specific mutated genes for which we identified homologs in *Chlamydomonas, Volvox,* and *Astrephomene* (Figure 3B-3E). We used existing datasets to determine whether these homologs had somatic-specific expression in *Volvox* (Matt and Umen 2017) or *Astrephomene* (Yamashita et al. 2021) or cold-induced expression in *Chlamydomonas* (L. Li et al. 2020). We identified homologs in all three species for 14 of the 18 annotated 11A5-specific genes and 24 of the 31 annotated 1A-specific genes. A significantly higher proportion of 11A5-specific mutated genes (10/14; 71.4%) had homologs with both cold-induced expression in *Chlamydomonas* and soma-specific expression in *Astrephomene* or *Volvox* than did 1A (7/24; 29.2%) (Fisher’s exact test, odds ratio = 5.75, 95% confidence interval = 1.17 – 34.50, p < 0.05) (Figure 3B-3D). In 11A5, 13 of the 14 genes (92.9%) had cold-induced or soma-specific expression, while only 16 of 24 (66.7%) in 1A did; however, this difference was not significant at the α = 0.05 level (Fisher’s exact test, odds ratio = 6.24, 95% confidence interval = 0.68 – 309.66, p > 0.05). Additionally, 3 11A5 genes had homologs with soma-specific expression in both *Volvox* and *Astrephomene* as well as cold-induced expression in *Chlamydomonas;* 1A had one such gene (Figure 3D). Additional details are provided in supplementary material S1, including supplementary table S17. The genes mutated only in 11A5 are therefore concentrated at the intersection of the cold shock response and the somatic differentiation pathways.

## Discussion

We report the experimental evolution of somatic differentiation in a predominantly undifferentiated species, demonstrating that changes in the regulation of an ancestrally plastic trait can rapidly shape a key step in an evolutionary transition. Modifications to somatic cell development following multiple rounds of cold shock occurred repeatedly across many independent populations with slightly varied selection protocols. In our CURE, 41.3% (19/46) of the student teams evolved lines with significant increases in the proportion of colonies with at least one somatic cell. Additionally, the proportion of colonies with soma increased in both of our lab-evolved lines. A subclone from one of those lines was consistently differentiated, showing that obligate differentiation can evolve in response to repeated stressors.

The proportion of somatic cells per differentiated colony converged with *Pleodorina,* a polyphyletic volvocine genus with naturally occurring obligate somatic differentiation (Figures 1 and 2). We found that the proportion of somatic cells per 16-celled differentiated colony increased relative to controls in 11 lines. In 3 of these lines, the proportion of somatic cells per differentiated 16-celled colony was significantly greater than that of 16-celled *P. starrii,* and consistent with the range reported for *P. californica.* In 8 of these lines, the proportion of somatic cells per 16-celled differentiated colony in these lines fell into the range of *P. starrii*, *P. californica,* or *P. japonica.* Across all lines, the proportion of colonies with soma was significantly correlated with the proportion of soma per differentiated colony (Figure 2B). We hypothesize that the two are mechanistically correlated, with somatic differentiation potentially being regulated by a threshold-dependent response, as proposed in Herron et al. (2014). The ease and rapidity with which a *Pleodorina*-like proportion of somatic cells evolved in *Eudorina* in response to a temporally fluctuating environment is consistent with the repeated evolution of the polyphyletic *Pleodorina* genus in nature.

We hypothesize that obligate differentiation in 11A5 evolved through modifications to stress-response genes that can also affect cell fate. The mutated genes in 11A5 predominantly lie in the intersection of the cold shock and somatic differentiation modules in volvocine homologs. Of the 11A5-specific mutated genes for which we identified homologs in *Chlamydomonas, Astrephomene,* and *Volvox,* 92.9 % (13/14) had homologs with soma-specific expression and/or cold-induced expression. 71.4% of these genes had homologs with both soma-specific and cold-induced expression, significantly more than in the control. We hypothesize that the stabilization of the plastic response occurred through changes in the same mechanisms that regulate the plastic response.

Co-option and genetic modification of stress-response modules may have shaped the parallel evolution of cellular differentiation in the volvocine algae. In *Volvox carteri,* the gene *regA* encodes a transcription factor necessary to maintain somatic differentiation (Kirk et al. 1999) and likely acts as a repressor. Somatic cells in *regA* mutants dedifferentiate and regain their reproductive potential. *RegA* belongs to a family of VARL domain genes called the reg cluster, which arose through gene duplication and descended from a gene likely similar to the *Chlamydomonas* stress-response gene *RLS1* (Duncan et al. 2007; Grochau-Wright et al. 2023) that suppresses cell division (Saggere et al. 2022). This stress-response gene was co-opted during the evolution of obligate somatic differentiation and is now developmentally regulated rather than exclusively environmentally regulated, though it retains the capacity to be upregulated in response to environmental stress (Olson and Nedelcu 2016; Nedelcu 2009; Nedelcu and Michod 2006; König and Nedelcu 2020).

The gene *regA* is not present in *Astrephomene* (Grochau-Wright et al. 2017); somatic differentiation is therefore regulated differently than in *Volvox.* Four transcription factors have been identified as candidate master regulators of *Astrephomene* somatic differentiation: two that belong to the MYB gene family, one that belongs to the RWP-RK gene family, and a homolog of reg cluster gene *rlsH* (Yamashita et al. 2021). In *Chlamydomonas,* an RWP-RK domain-containing protein is a hub gene regulating the co-expression network of the cold stress response, and the MYB-related family of transcription factors has the second-highest number of differentially expressed genes during one hour of cold shock (L. Li et al. 2020).

We did not identify 11A5-specific mutations in any reg cluster, MYB, or RWP-RK genes. However, we found that gene EUDS_000603, which likely encodes a transcription factor with SBP and BTB/POZ domains, had a multi-exon deletion in 11A5 that likely caused the DNA-binding domain to be lost. Squamosa binding-like proteins are developmental regulators that affect the transition from vegetative to reproductive states (X. Chen et al. 2010); a key characteristic of somatic cells is their lack of reproduction (Nozaki et al. 1989). The multiple genetic and developmental pathways to somatic differentiation available to volvocine algae lineages may help explain the parallel evolution of soma in this clade. While *Astrephomene, Volvox*, and 11A5 likely utilize different mechanisms regulating differentiation, they all may have co-opted components of stress-response modules during the transition to cellular differentiation.

Regardless of the mechanisms involved, the evolution of somatic differentiation underpins the evolution of a new kind of evolutionary individual: an integrated multicellular organism with reproductive division of labor. Somatic cells do not reproduce and instead specialize on the survival of the multicellular group. Consequently, specialized cells would have low fitness outside of the context of the multicellular group. Once differentiation evolves, cell fitness depends on the fitness of the multicellular organism, and selection operates on multicellular organisms rather than on cells. This is therefore the first study to report the experimental evolution of a new kind of evolutionary individual with reproductive division of labor.

We found that changes in plasticity can mediate the evolution of a new kind of evolutionary individual. The environmental stress that induces the plastic phenotype likely also selected for changes in its regulation, as we did not directly select for differentiation. Consistent with this possibility, modeling work has shown that somatic specialization evolves in response to repeated stressors when somatic cells provide survival benefits (C. Zhang et al. 2026) Genetic variation arose during the experiment, as we started all populations from a single clonally reproducing colony and found that not all lab line subclones produced a mix of differentiated and undifferentiated offspring, since 11A5 consistently developed somatic cells. In 11A5, numerous changes occurred in genes that are likely to affect the response to cold shock, the stressor that induces plastic development of soma in *Eudorina.* We hypothesize that the repeatability of the evolution of somatic differentiation comes from alignment between plastic responses, selection pressures, and the generation of genetic variation. Such an alignment may increase the evolvability of the plastic response and allow for processes such as genetic assimilation, the genetic stabilization of a plastic response, to proceed rapidly.

### Limitations and future work

Additional research is needed to better understand this transition. First, we don’t understand how selection shaped this transition; we need fitness data to determine whether and how plastic differentiation protects against cold shock. Second, we only sequenced a single experimentally evolved strain, and therefore, we don’t know if other strains have changes in different genes. We may also have missed genes important for differentiation during alignment, mutation identification, and filtering. Of the sample-specific mutated genes we identified, it’s unclear which genetic changes directly cause the differentiated phenotype and how many of the identified genetic changes have functional consequences. Any given sample-specific mutation may have contributed to adaptation to cold shock, been part of a response to genetic changes that occurred during repeated cold shock, been selected for by culture collection conditions, or changed in frequency in either line due to genetic drift. Identified sample-specific mutations could be neutral or could be sequencing artifacts. Sequencing additional experimentally evolved strains is necessary to better understand the genetic architecture that underpins somatic differentiation in *Eudorina.* Finally, transcriptomic data are needed to test whether the expression of the cold shock response pathway has been modified and to compare the cold shock response in *Eudorina* to the experimentally evolved changes in gene expression that occurred due to repeated exposure to cold stress.

### Conclusions

Somatic differentiation is a key step in the transition to multicellular individuality. We found that it can evolve rapidly and repeatedly through the modification of plastic responses to the environment. Repeated exposure to cold shock led to an increased proportion of colonies with soma in 44% of lines (21/48), as well as an increased proportion of soma per differentiated colony in 23% of lines (11/48). These 11 lines converged on the proportion of somatic cells per differentiated colony that is seen in multiple *Pleodorina* species. We also found that plastic differentiation became obligate in one lineage in which all 16-celled colonies had somatic cells; we identified this lineage more than 50 generations after the last round of cold shock. Genes with mutations in this lineage were concentrated at the intersection of the cold-shock response and somatic differentiation pathways, suggesting that the stabilization of somatic development may have been shaped by stress-response genes whose homologs can affect cell fate in other species. This is the first published example of the experimental evolution of obligate somatic differentiation, and it shows that environmentally induced developmental changes can mediate a major evolutionary transition.

## Acknowledgements

We thank all members of the Michod and Olson labs, with special thanks to Aurora Nedelcu, SoRi La, De Andre Onyi, Zach Grochau-Wright, Arthur Cousson, Lidia Lopez, Lily Alberding, and Lucas Williams for their assistance with past and ongoing plasticity and experimental evolution research. We thank Matthew Herron, Fred Nijhout, Mike Travisano, William Ratcliff, Beatriz Baselga-Cervera, Aurora Nedelcu, and Nicholas Levis for productive conversations and feedback. We thank Hisayoshi Nozaki for providing *Eudorina* sp. NIES 3984 to the Michod Lab, SoRi La for subcloning *Eudorina* for the ECOL 182L CURE, and Matthew Herron for providing the raw data associated with Herron et al. (2014). We would especially like to thank the ECOL 182L laboratory coordinator, Oona Husok, for maintaining the CURE populations and managing CURE logistics. We thank the ECOL 182L teaching assistants and all the students who participated in the course. Full acknowledgments associated with the CURE, including 256 undergraduate students who participated and consented to be recognized by name, are given in supplementary material S2. This work was funded by NSF grant DEB-2029999 and is based upon work supported by the NSF Postdoctoral Research Fellowships in Biology Program under grant no. 2209373.

