## Supplementary materials S1 for "Somatic differentiation evolves rapidly and repeatedly through the modification of developmental plasticity"

#### Supplementary results S1

##### Experimental evolution: lab lines

**S1.** The proportion of colonies with somatic cells for each of the four lab lines, with the frequency of each flask shown separately.

| Treatment | Line | Replicate flask | Number of colonies | Number of differentiated colonies | Frequency of differentiation |
| --- | --- | --- | --- | --- | --- |
| Control | 1A | 1 | 100 | 10 | 0.10 |
| Control | 1A | 2 | 102 | 20 | 0.20 |
| Control | 9A | 1 | 101 | 9 | 0.090 |
| Control | 9A | 2 | 101 | 20 | 0.20 |
| Cold | 11A | 1 | 102 | 48 | 0.47 |
| Cold | 11A | 2 | 101 | 42 | 0.42 |
| Cold | 19A | 1 | 100 | 42 | 0.42 |
| Cold | 19A | 2 | 102 | 42 | 0.41 |

**Table S2.** Proportion of somatic cells per differentiated colony for each of the four lab lines.

| Treatment | Line | N | Mean proportion of soma | Median proportion of soma | Mean number of soma | Percent single somatic cell |
| --- | --- | --- | --- | --- | --- | --- |
| Control | 1A | 30 | 0.13 | 0.063 | 2.00 | 60.0% |
| Control | 9A | 29 | 0.18 | 0.13 | 2.79 | 48.3% |
| Cold | 11A | 90 | 0.24 | 0.16 | 3.62 | 34.4% |
| Cold | 19A | 84 | 0.22 | 0.19 | 3.18 | 34.5% |
| Cold | 11A5 | 64 | 0.41 | 0.34 | 5.50 | 23.4% |

**Table S3.** Proportion of somatic cells per differentiated colony for the two treatments, with lines pooled.

| Treatment | Lines | N | Mean proportion of soma | Median proportion of soma | Q25 – Q75 | Min - max |
| --- | --- | --- | --- | --- | --- | --- |
| Control | 1A and 9A | 59 | 0.16 | 0.13 | 0.063 – 0.13 | 0.063 – 0.81 |
| Cold | 11A and 19A | 174 | 0.23 | 0.19 | 0.063 – 0.31 | 0.063 – 0.88 |

**Table S4.** Proportion of somatic cells per differentiated colony for 11A5 and 1A. Data from 1A is taken from the direct comparison with 11A5. 11A5 had a significantly higher proportion of somatic cells per differentiated colony than did controls (Mann-Whitney U test, $W = 32$, $p < 0.0001$).

| Treatment | Line | N | Mean proportion of soma | Median proportion of soma | Q25 – Q75 | Min - max |
| --- | --- | --- | --- | --- | --- | --- |
| Control | 1A | 9 | 0.08 | 0.063 | 0.063 – 0.063 | 0.063 – 0.13 |
| Cold | 11A5 | 64 | 0.41 | 0.34 | 0.19 – 0.64 | 0.031 – 0.94 |

###### Experimental evolution: CURE

The proportion of differentiated colonies in a line predicted the proportion of soma per differentiated colony in that line (odds ratio = 1.11 per 10-percentage-point increase, 95% confidence interval = 1.06 – 1.17, $p < 0.0001$). This relationship did not differ between control and experimentally evolved lines (interaction likelihood-ratio $\chi^2_1 = 0.49$, $p > 0.05$). When the proportion of differentiated colonies was included in the analysis, the experimental treatment did not explain additional variation in the proportion of somatic cells per differentiated colony ($\chi^2_1 = 0.03$, $p > 0.05$). The proportion of differentiated colonies remained a strong predictor of the proportion of somatic cells per differentiated colony after accounting for experimental treatment ($\chi^2_1 = 15.5$, $p < 0.0001$), and accounted for approximately 37% of the variance among lines. Model selection by AIC favored the proportion of differentiated colonies as a predictor of the proportion of soma over any model that included experimental treatment. The relationship between proportion of colonies with soma and proportion of soma per differentiated colony did not differ between experimentally evolved lines in which the proportion of colonies with soma evolved and lines in which it did not (interaction likelihood-ratio $\chi^2_1 = 0.40$, $p > 0.05$).

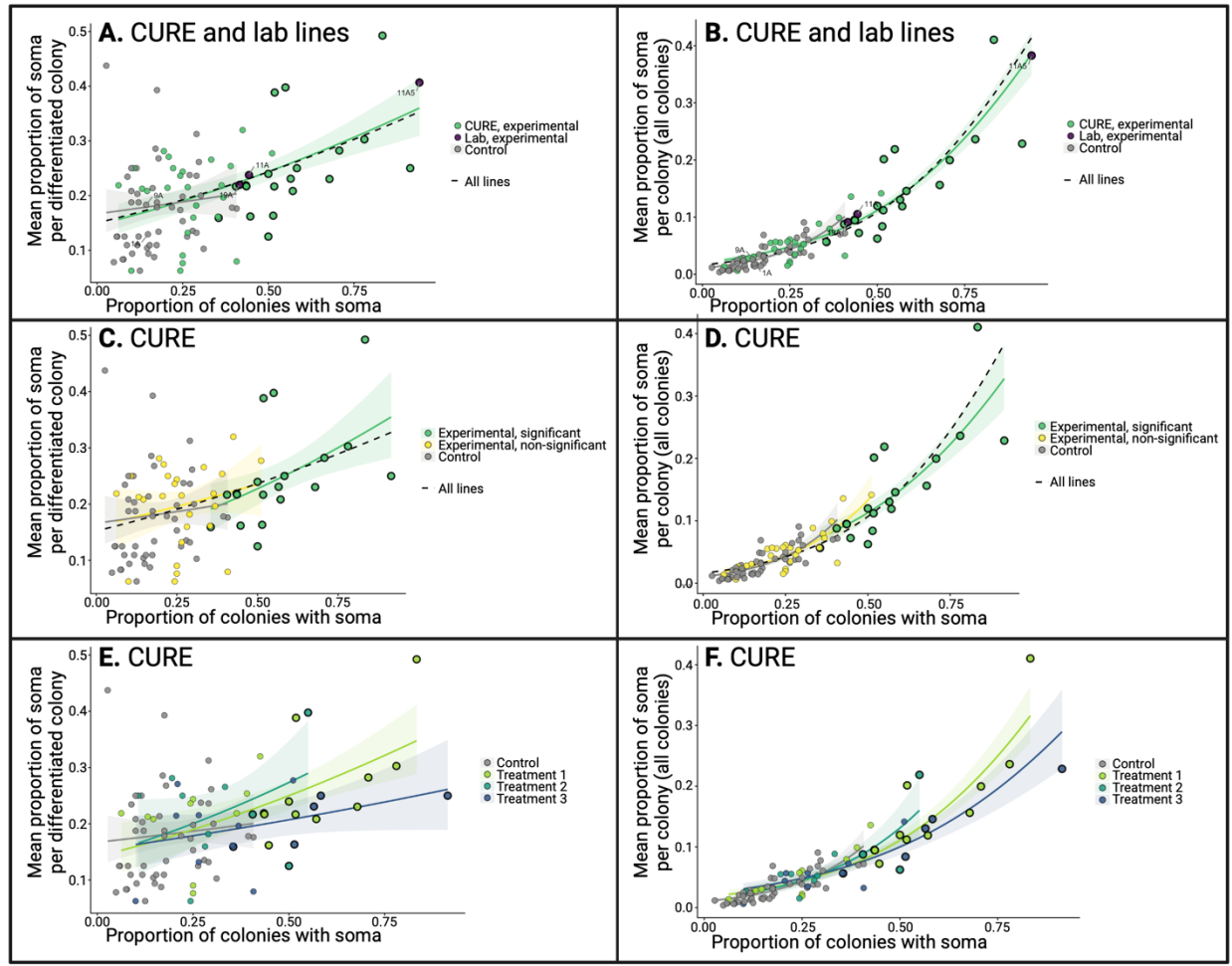

**Figure S1.** Relationship between the proportion of colonies with soma and the proportion of soma per differentiated colony (A, C, and E) or per colony (B, D, and F). In panels B, D, and F, the relationship between the proportion of colonies with soma and the mean proportion of soma per colony (for all colonies) is automatically positively correlated because the proportion of colonies with soma affects the mean proportion of soma per colony when undifferentiated colonies are included. Each point is a single laboratory or CURE line. Lines in which the proportion of colonies with somatic cells increased significantly are shown with a bold outline. Controls are shown in gray in all panels. Colored lines are fitted values from a beta-binomial generalized linear mixed model, with line as a random effect and group-specific slopes. The shaded bands are 95% confidence intervals. A-B) Both CURE and lab lines are shown. The green dots are experimentally evolved lines from the CURE, and the purple dots are experimentally evolved lab lines. The green line and 95% confidence interval are derived from both the CURE and lab lines. The gray line and 95% confidence interval are derived from the pooled CURE and lab line controls. The dashed black line corresponds to a single pooled slope fitted across all lines shown in the panel. C-D) Only CURE lines are shown. The green dots, line, and 95% confidence interval are experimentally evolved lines that have significant changes in the proportion of colonies with soma. The yellow dots, line, and 95% confidence interval are experimentally evolved lines that do not have significant changes in the proportion of colonies

with soma. The gray dots, line, and 95% confidence interval are controls. The dashed black line corresponds to a single pooled slope fitted across all lines shown in the panel. E-F). The three CURE treatments are shown. Light green dots, line, and 95% confidence interval are experimentally evolved lines from treatment 1. Turquoise dots, line, and 95% confidence interval are experimentally evolved lines from treatment 2. Dark blue dots, line, and 95% confidence interval are experimentally evolved lines from treatment 3. Gray dots, line, and 95% confidence interval are controls. Figure created in https://BioRender.com

**Table S5.** Proportion of colonies with somatic cells, experimental vs. control treatments.

| Model | Log odds ratio | Standard error | Odds ratio | 95% confidence interval | P-value |
| --- | --- | --- | --- | --- | --- |
| GLM (colonies independent) | 1.08 | 0.083 | 2.95 | 2.51 – 3.47 | <0.0001 |
| GEE, exchangeable (cluster = line) | 1.09 | 0.15 | 2.98 | 2.21 – 4.03 | <0.0001 |
| GLMM, binomial (cluster = line) | 1.16 | 0.17 | 3.18 | 2.28 – 4.44 | <0.0001 |

**Table S6.** Line-level proportion of differentiated colonies for pooled control and experimentally evolved lines.

|  | Control | Cold shock |
| --- | --- | --- |
| Number of lines | 46 | 46 |
| Total number of colonies | 1,608 | 1,602 |
| Differentiated colonies | 281 | 616 |
| Frequency of differentiation | 0.18 | 0.39 |
| Median number of colonies per line | 36 | 36 |
| Mean frequency of differentiation per line (SD) | 0.18 (0.093) | 0.39 (0.20) |
| Median frequency of differentiation per line (SD) | 0.16 | 0.37 |
| Range | 0.027-0.41 | 0.063-0.91 |

**Table S7.** Interaction tests of treatment (cold shock vs. control) x treatment protocol (treatments 1 (N = 21), 2 (N = 10), and 3 (N = 15)).

| Test | Statistic | df | P-value |
| --- | --- | --- | --- |
| GEE, Wald; interaction (treatment x protocol) | $\chi^2 = 1.78$ | 2 | >0.05 |
| GLMM, likelihood ratio; interaction (treatment x protocol) | $\chi^2 = 1.32$ | 2 | >0.05 |

**Table S8.** The number and proportion of differentiated colonies for pooled control and experimentally evolved lines from each treatment protocol.

| Treatment protocol | Treatment | N lines | N colonies | N differentiated | Pooled proportion differentiated | Line-level mean proportion differentiated (SD) |
| --- | --- | --- | --- | --- | --- | --- |
| Tx 1 | Control | 21 | 732 | 126 | 0.17 | 0.18 (0.11) |
| Tx 1 | Cold shock | 21 | 748 | 315 | 0.42 | 0.43 (0.21) |
| Tx 2 | Control | 10 | 350 | 60 | 0.17 | 0.17 (0.082) |
| Tx 2 | Cold shock | 10 | 339 | 104 | 0.31 | 0.31 (0.14) |
| Tx 3 | Control | 15 | 526 | 95 | 0.18 | 0.18 (0.086) |
| Tx 3 | Cold shock | 15 | 515 | 197 | 0.38 | 0.39 (0.20) |

**Table S9.** The proportion of somatic cells per differentiated colony, comparing control vs. cold-shocked colonies and clustering by line.

| Model | Log odds ratio | Standard error | Odds ratio | 95% confidence interval | P-value |
| --- | --- | --- | --- | --- | --- |
| GEE, exchangeable | 0.29 | 0.11 | 1.33 | 1.08-1.64 | <0.01 |
| GLMM, binomial | 0.29 | 0.11 | 1.33 | 1.08-1.65 | <0.01 |
| GLMM, beta-binomial | 0.26 | 0.10 | 1.29 | 1.07-1.56 | <0.01 |

**Table S10.** The proportion of somatic cells per differentiated colony for all control and all experimentally evolved lines, lines with significant changes in the proportion of colonies with soma and their corresponding controls, and lines without significant changes in the proportion of colonies with soma and their corresponding controls.

| Lines | Treatment | N colonies | N lines | Mean | Median |
| --- | --- | --- | --- | --- | --- |
| All | Control | 281 | 46 | 0.18 | 0.125 |
| All | Cold shock | 616 | 46 | 0.24 | 0.19 |
| Significant changes in the proportion of colonies with soma | Control | 100 | 19 | 0.16 | 0.125 |
| Significant changes in the proportion of colonies with soma | Cold shock | 352 | 19 | 0.27 | 0.19 |
| No significant changes in the proportion of colonies with soma | Control | 181 | 27 | 0.19 | 0.125 |
| No significant changes in the proportion of colonies with soma | Cold shock | 264 | 27 | 0.21 | 0.125 |

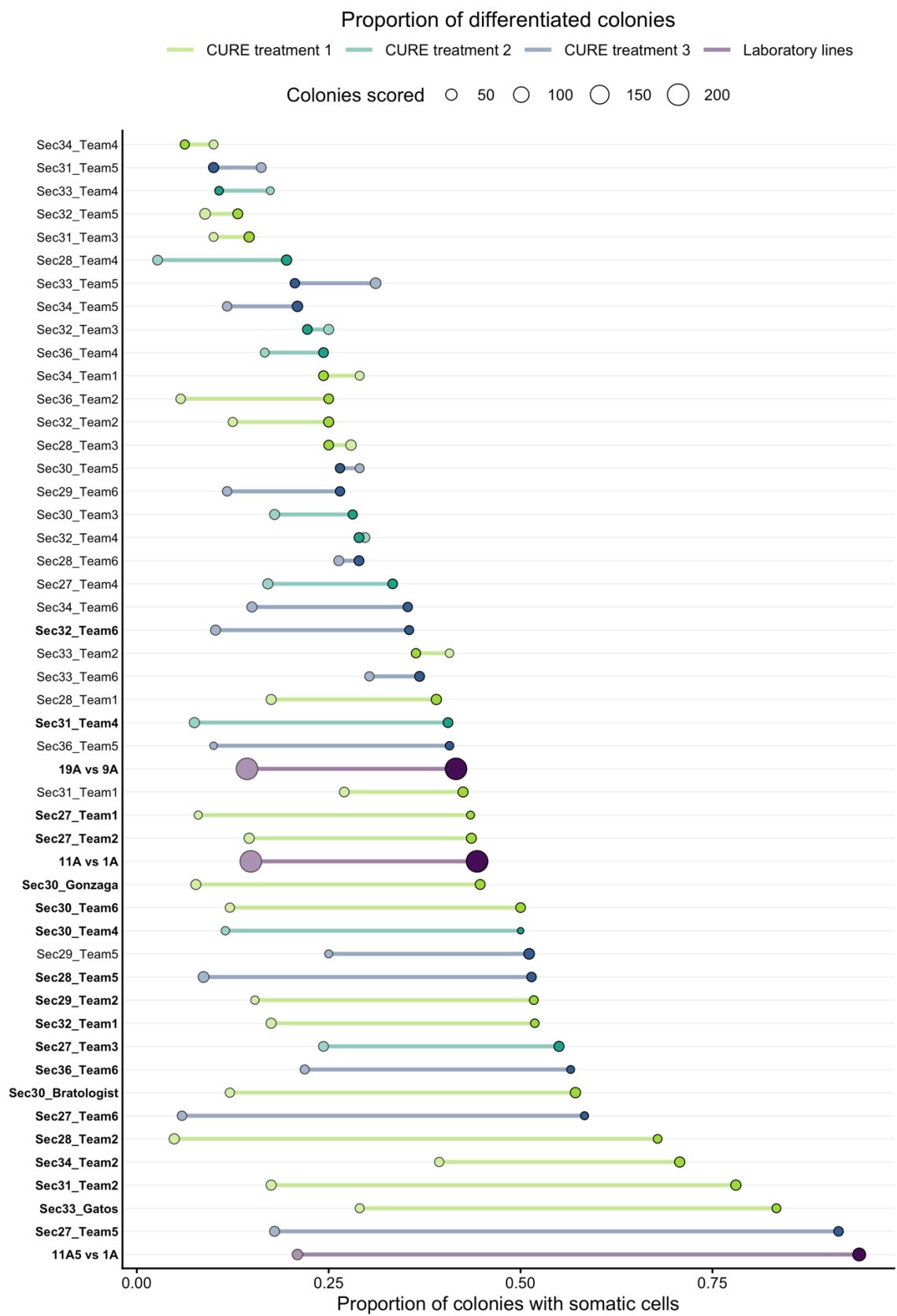

**Figure S2.** Proportion of colonies with somatic cells in each experimentally evolved line and its paired control. Each row is one experimentally evolved line joined to its paired control by a horizontal line. Filled points are experimentally evolved lines, and pale points are the paired control. Point size corresponds to the number of colonies scored. Color denotes the treatment: light green is CURE treatment 1, turquoise is CURE treatment 2, dark blue is CURE treatment 3, and purple is laboratory. Rows are ordered by the proportion of colonies with soma for each experimentally evolved line. Names correspond to the name of each CURE team or to the lab line comparison. Names in bold correspond to experimentally evolved lines in which the proportion of colonies with soma is significantly different from the paired control.

### Somatic proportion, differentiated colonies only

CURE treatment 1 CURE treatment 2 CURE treatment 3 Laboratory lines

Differentiated colonies ○ 20 ○ 40 ○ 60 ○ 80

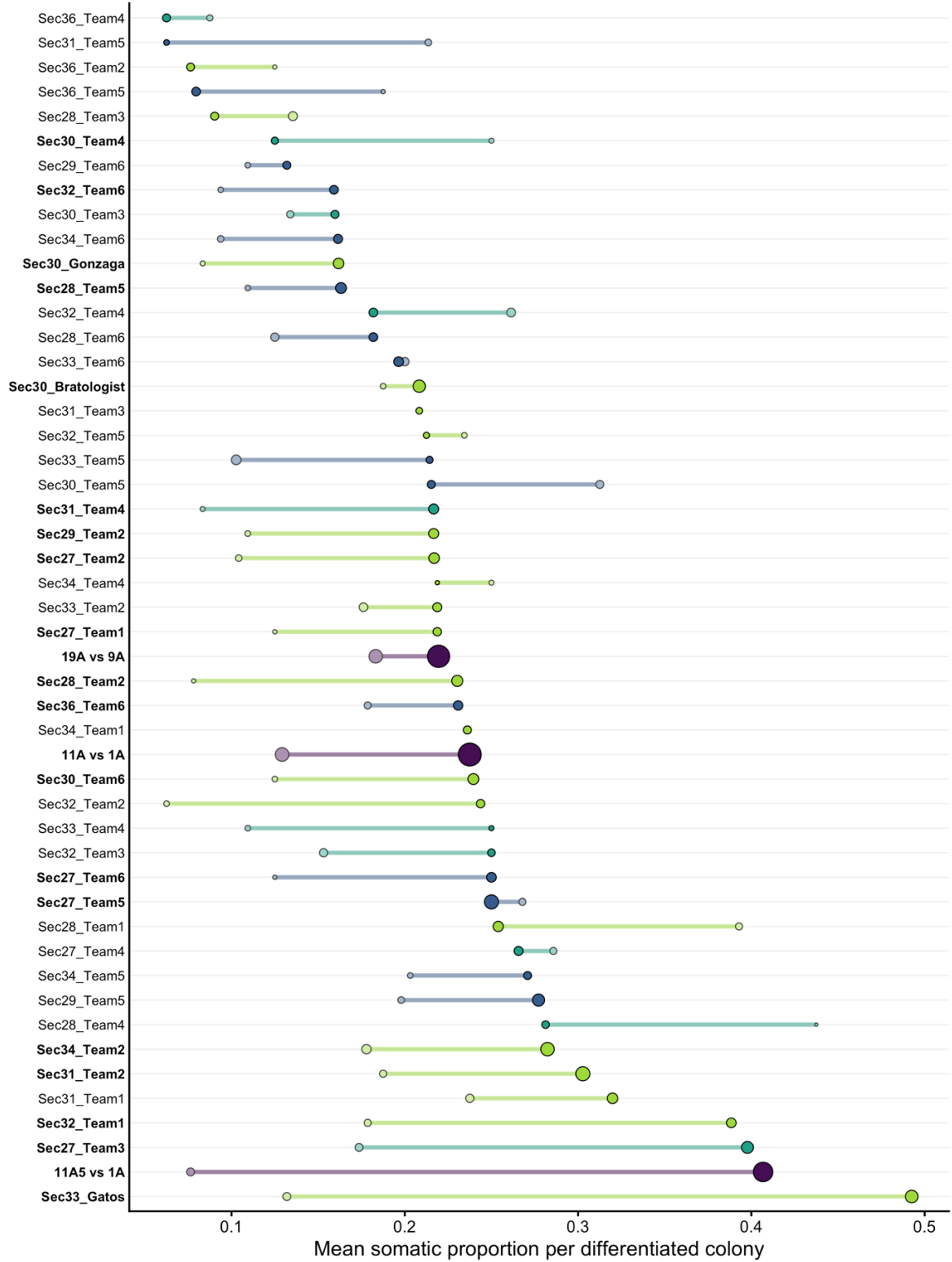

**Figure S3.** Proportion of somatic cells per differentiated colony in each experimentally evolved line and its paired control. Each row is one experimentally evolved line joined to its paired control by a horizontal line. Filled points are experimentally evolved lines, and pale points are the paired control. Point size corresponds to the number of colonies scored. Color denotes the treatment: light green is CURE treatment 1, turquoise is CURE treatment 2, dark blue is CURE treatment 3, and purple is laboratory. Rows are ordered by the proportion of somatic cells per differentiated colony for each experimentally evolved line. Names correspond to the name of each CURE team or to the lab line comparison. Names in bold correspond to experimentally evolved lines in which the proportion of colonies with soma is significantly different from the paired control.

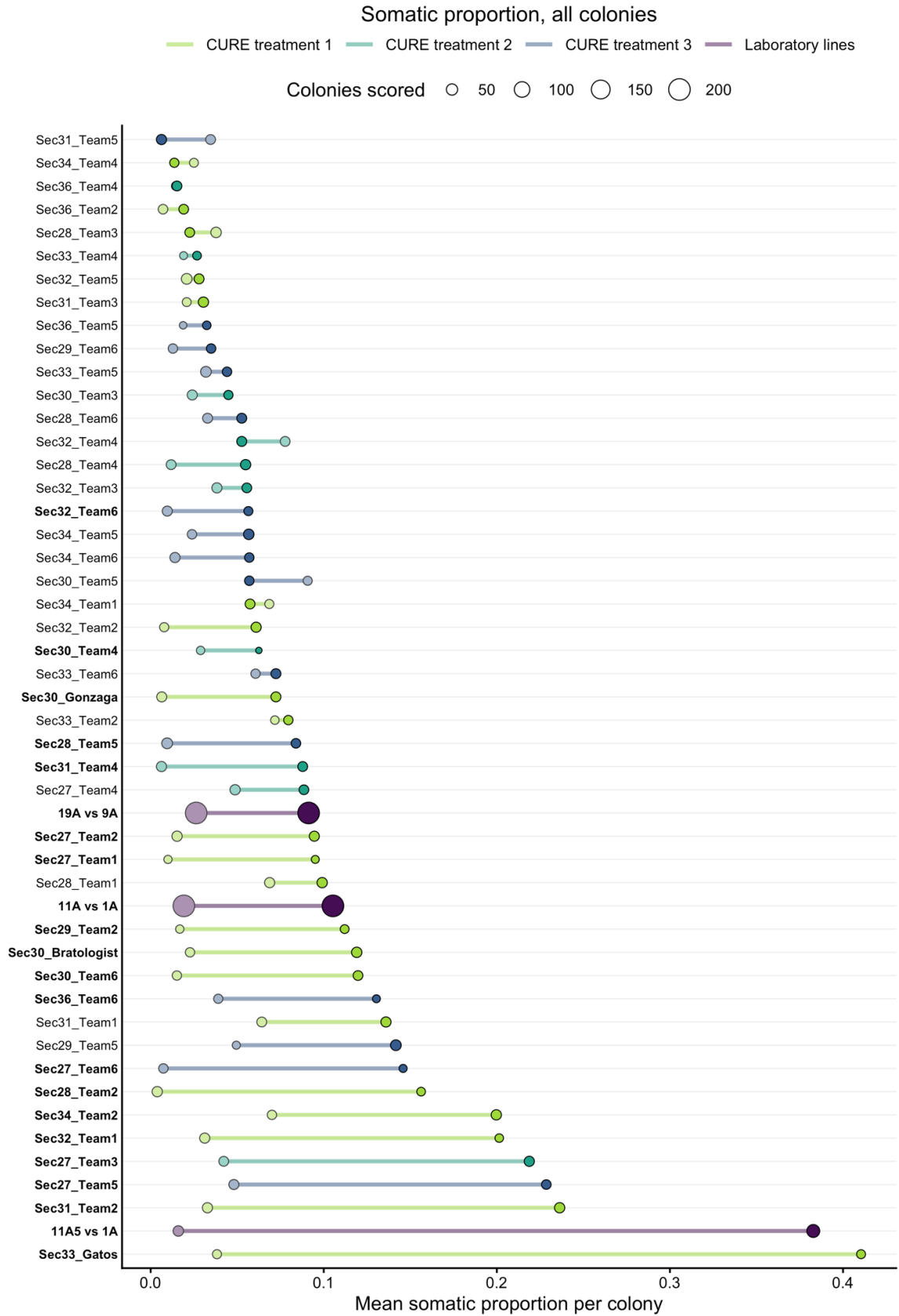

**Figure S4.** Proportion of somatic cells per colony (including both differentiated and undifferentiated colonies) in each experimentally evolved line and its paired control. Each row is one experimentally evolved line joined to its paired control by a horizontal line. Filled points are experimentally evolved lines, and pale points are the paired control. Point size corresponds to the number of colonies scored. Color denotes the treatment: light green is CURE treatment 1, turquoise is CURE treatment 2, dark blue is CURE treatment 3, and purple is laboratory. Rows are ordered by the proportion of somatic cells per colony for each experimentally evolved line. Names correspond to the name of each CURE team or to the lab line comparison. Names in bold correspond to experimentally evolved lines in which the proportion of colonies with soma is significantly different from the paired control.

##### Comparison to *Pleodorina*

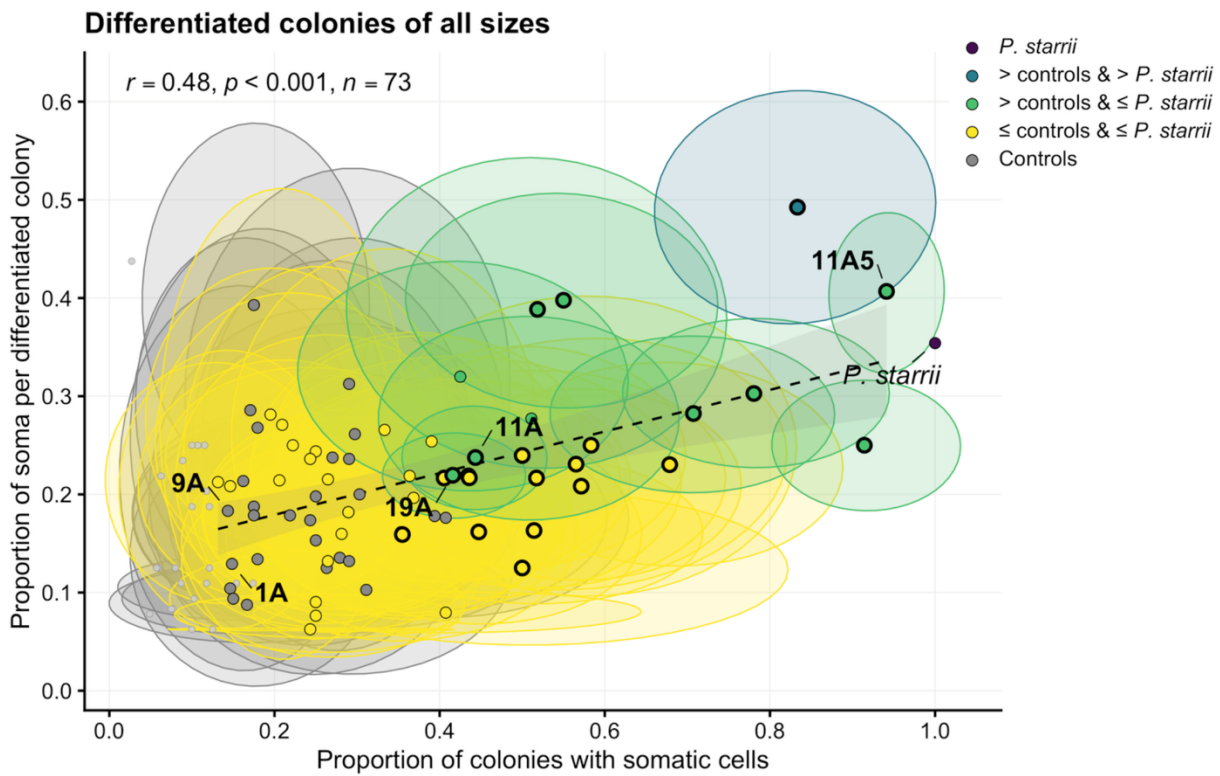

**Figure S5.** The proportion of differentiated colonies of all sizes plotted against the proportion of somatic cells per differentiated colony. Each line is represented by a single data point with a 95% confidence interval ellipse. *P. starrii* is shown in dark purple. The line for which the proportion of somatic cells per differentiated colony is significantly higher than *P. starrii* is shown in teal. The lines for which the proportion of somatic cells per differentiated colony is greater than the pooled controls but not greater than *P. starrii* are shown in green. Yellow corresponds to lines for which the proportion of somatic cells per differentiated colony is not greater than controls. Controls are shown in gray. Lines for which there are fewer than 5 differentiated colonies are represented as small, light gray points. The dashed line shows the relationship between the

proportion of colonies with somatic cells and the proportion of soma per differentiated colonies for all *Eudorina* lines with at least 5 differentiated colonies ($r = 0.52$, $p < 0.001$, $n = 73$).

##### Experimentally evolved lines

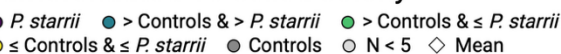

#### Control lines

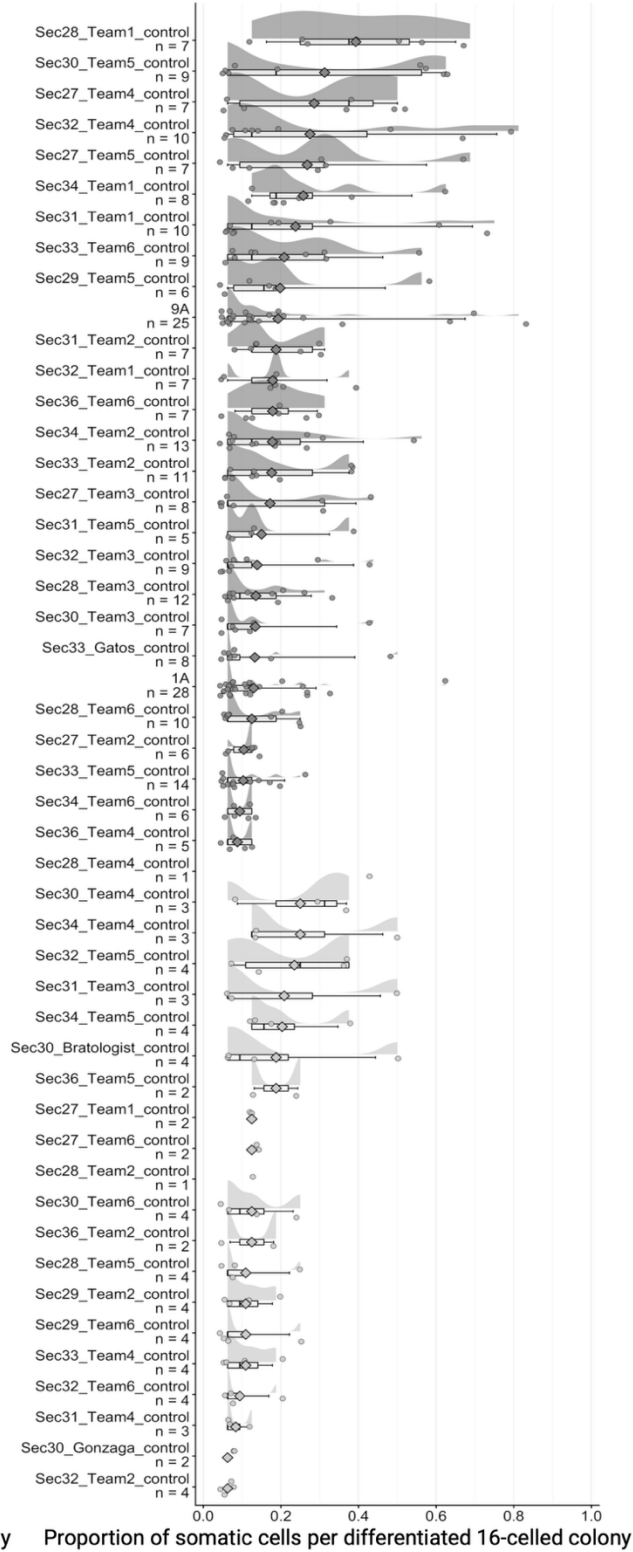

**Figure S6.** The distribution of the proportion of somatic cells per line for differentiated 16-celled colonies from all lines. The distribution of somatic cells for 16-celled *P. starrii* is shown at the top left in dark purple. The mean proportion of somatic cells for 16-celled *P. starrii* is shown in the dashed purple line. The experimentally evolved lines shown in teal are those in which the mean proportion of somatic cells per differentiated 16-celled colony is significantly greater than that of *P. starrii*. The mean number of somatic cells per differentiated 16-celled colony for each line is shown as a diamond. The lines shown in green are those for which the mean proportion of somatic cells per differentiated 16-celled colony is significantly greater than in pooled controls but not significantly greater than *P. starrii*. Experimentally evolved lines shown in yellow are those for which the proportion of somatic cells per differentiated 16-celled colony is not significantly greater than controls. Controls are shown in gray on the right. Lines with fewer than 5 data points are shown in light gray. *Eudorina* lines are ordered by the mean proportion of somatic cells per differentiated 16-celled colony. Created in https://BioRender.com.

#### Proportion of somatic cells per differentiated colony

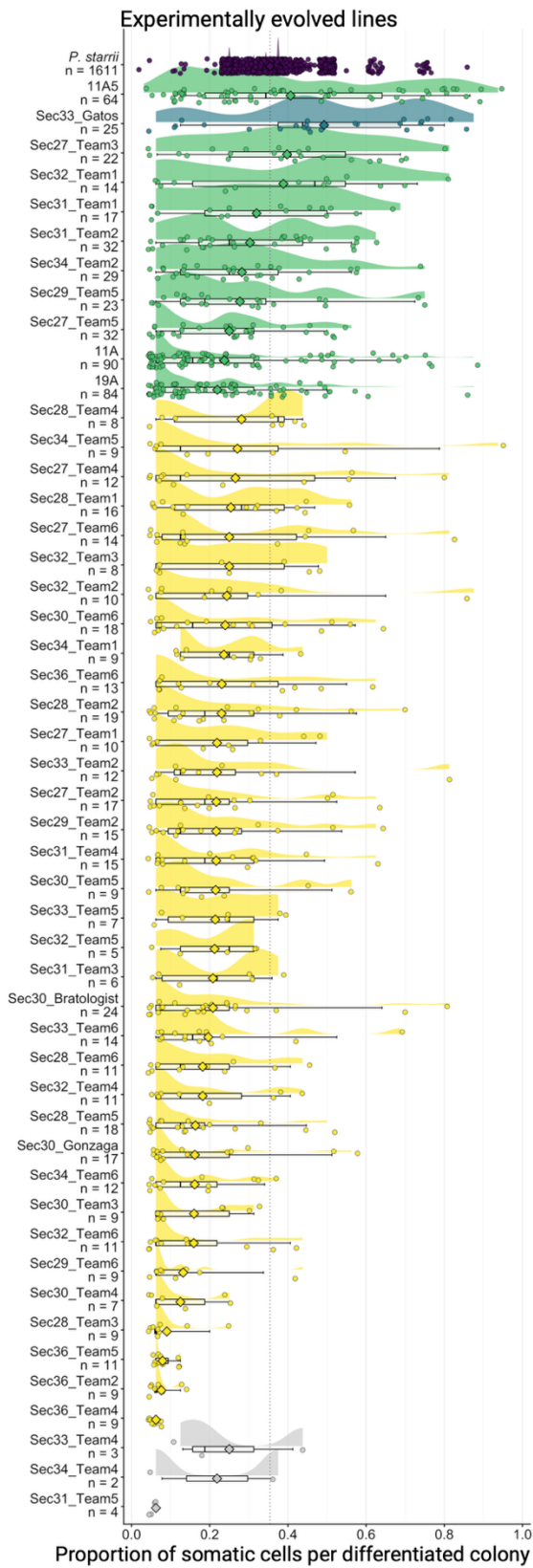

● *P. starrii* ● > Controls & > *P. starrii* ● > Controls & ≤ *P. starrii*
● ≤ Controls & ≤ *P. starrii* ● Controls ○ N < 5 ◇ Mean

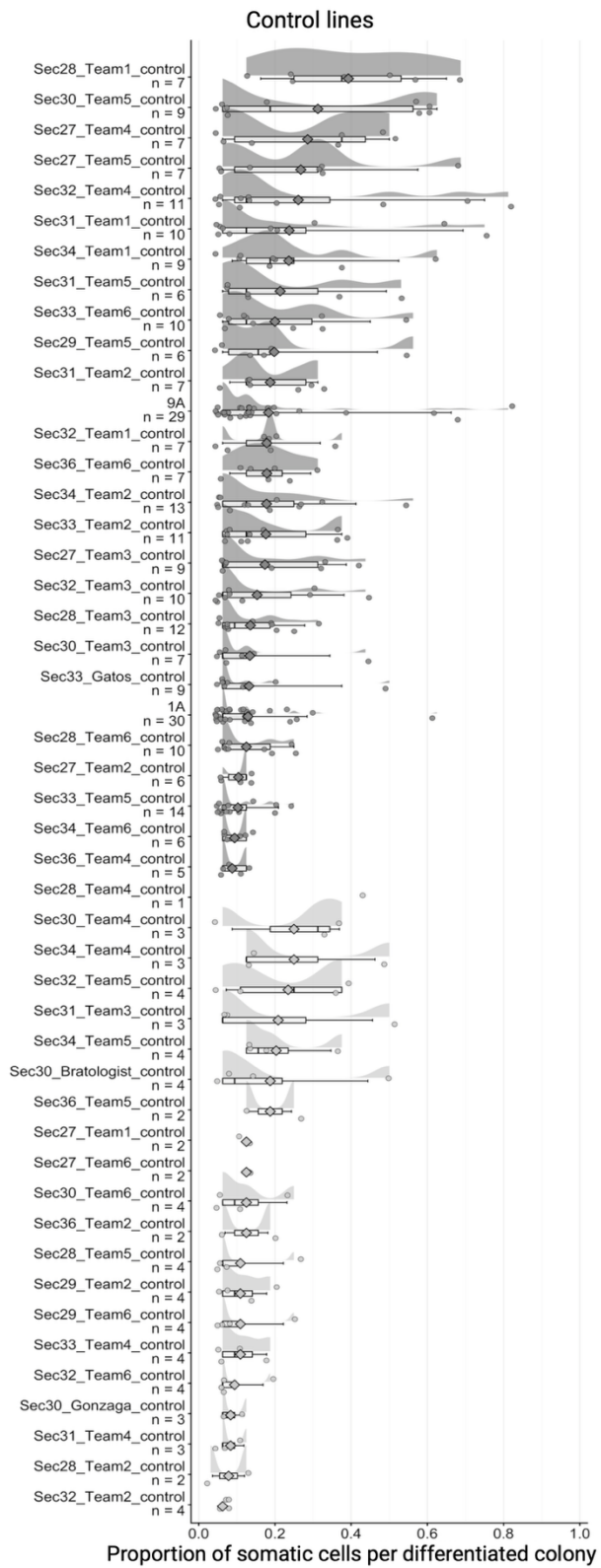

**Figure S7.** The distribution of the proportion of somatic cells per line for differentiated colonies of all sizes from all lines. The distribution of somatic cells for *P. starrii* is shown at the top left in dark purple. The mean proportion of somatic cells for *P. starrii* is shown in the dashed purple line. The experimentally evolved lines shown in teal are those in which the mean proportion of somatic cells per differentiated colony is significantly greater than that of *P. starrii*. The mean proportion of somatic cells per differentiated colony for each line is shown as a diamond. The lines shown in green are those for which the mean proportion of somatic cells per differentiated colony is significantly greater than in pooled controls but not significantly greater than *P. starrii*. Experimentally evolved lines shown in yellow are those for which the proportion of somatic cells per differentiated colony is not significantly greater than controls. Controls are shown in gray on the right. Lines with fewer than 5 data points are shown in light gray. *Eudorina* lines are ordered by the mean proportion of somatic cells per differentiated colony. Created in https://BioRender.com.

#### Proportion of somatic cells per 16-celled colony

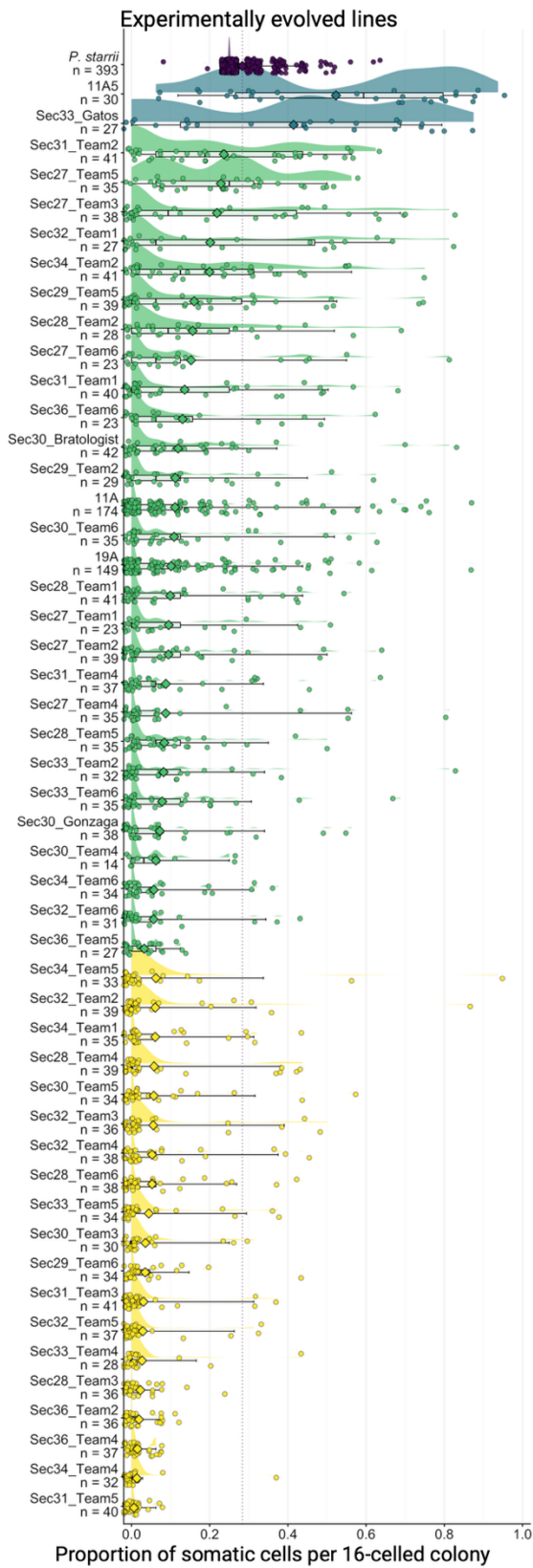

● *P. starrii* ● > Controls & > *P. starrii* ● > Controls & ≤ *P. starrii*
● ≤ Controls & ≤ *P. starrii* ● Controls ○ N < 5 ◇ Mean

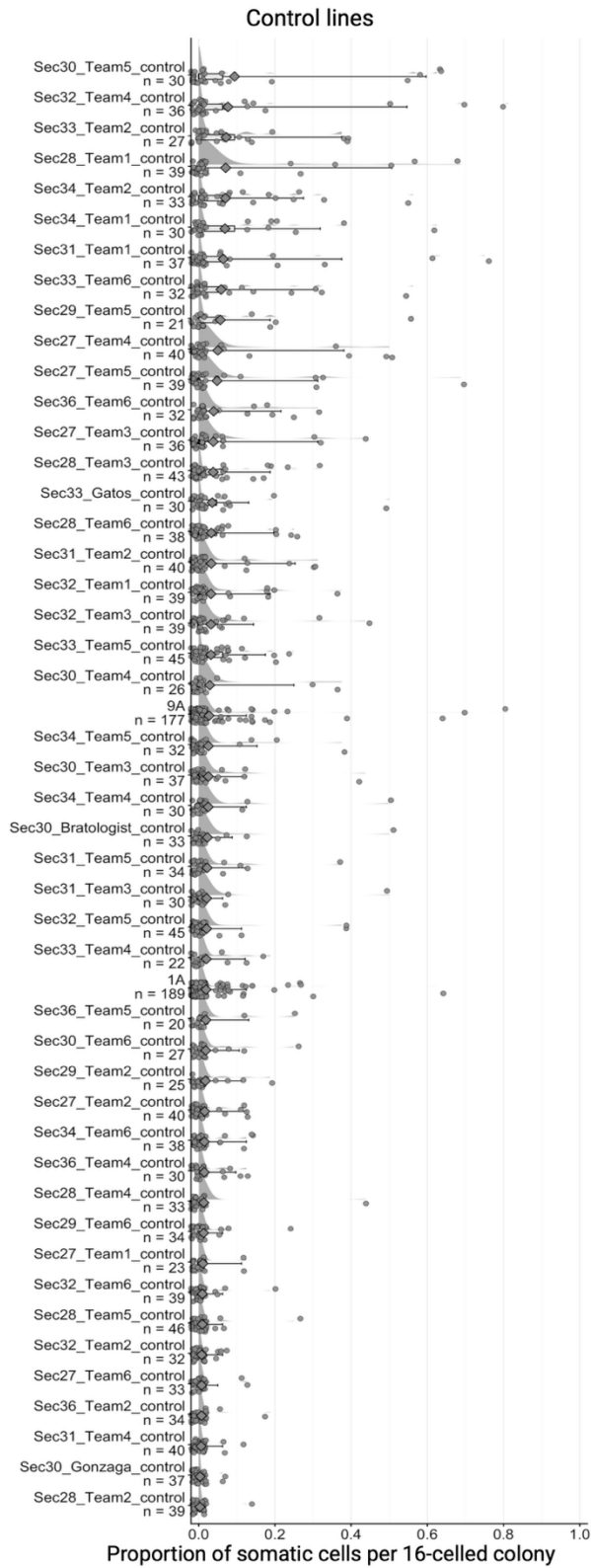

**Figure S8.** The distribution of the proportion of somatic cells per line for 16-celled colonies from all lines. The distribution of somatic cells for 16-celled *P. starrii* is shown at the top left in dark purple. The mean proportion of somatic cells for 16-celled *P. starrii* is shown in the dashed purple line. The experimentally evolved lines shown in teal are those in which the mean proportion of somatic cells per 16-celled colony is significantly greater than that of *P. starrii*. The mean number of somatic cells per 16-celled colony for each line is shown as a diamond. The lines shown in green are those for which the mean proportion of somatic cells per 16-celled colony is significantly greater than in pooled controls but not significantly greater than *P. starrii*. Experimentally evolved lines shown in yellow are those for which the proportion of somatic cells per 16-celled colony is not significantly greater than controls. Controls are shown in gray on the right. Lines with fewer than 5 data points are shown in light gray. *Eudorina* lines are ordered by the mean proportion of somatic cells per 16-celled colony. Created in https://BioRender.com.

#### Proportion of somatic cells per colony

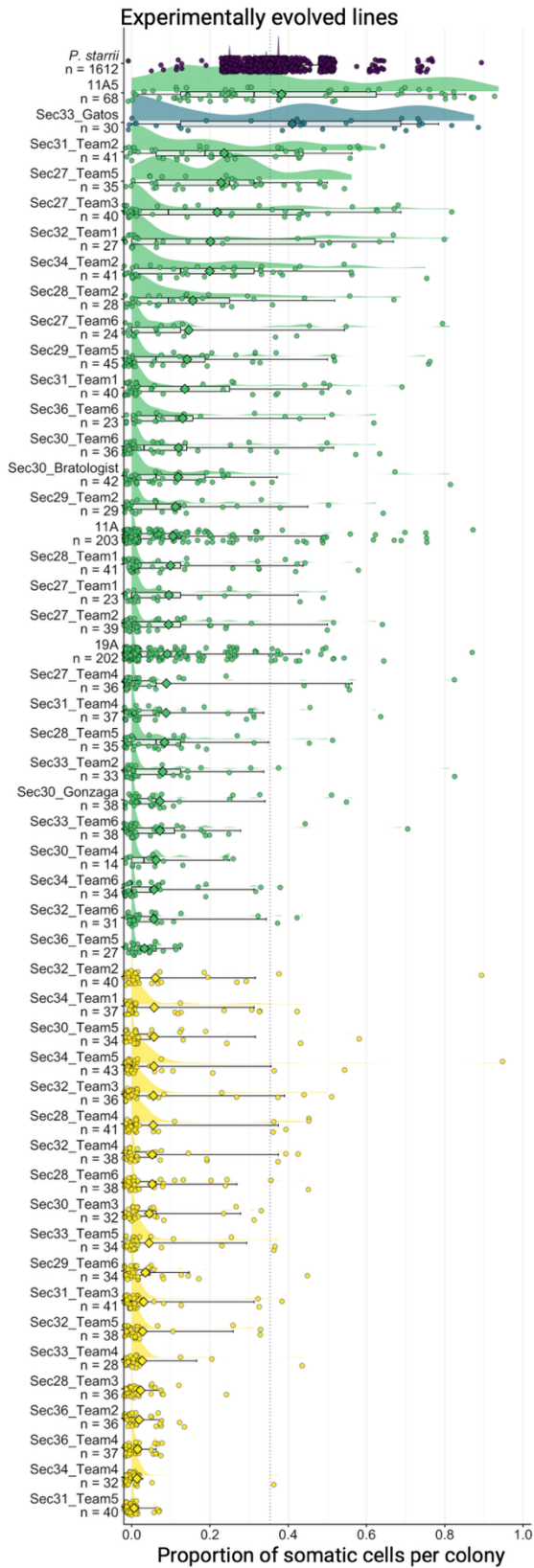

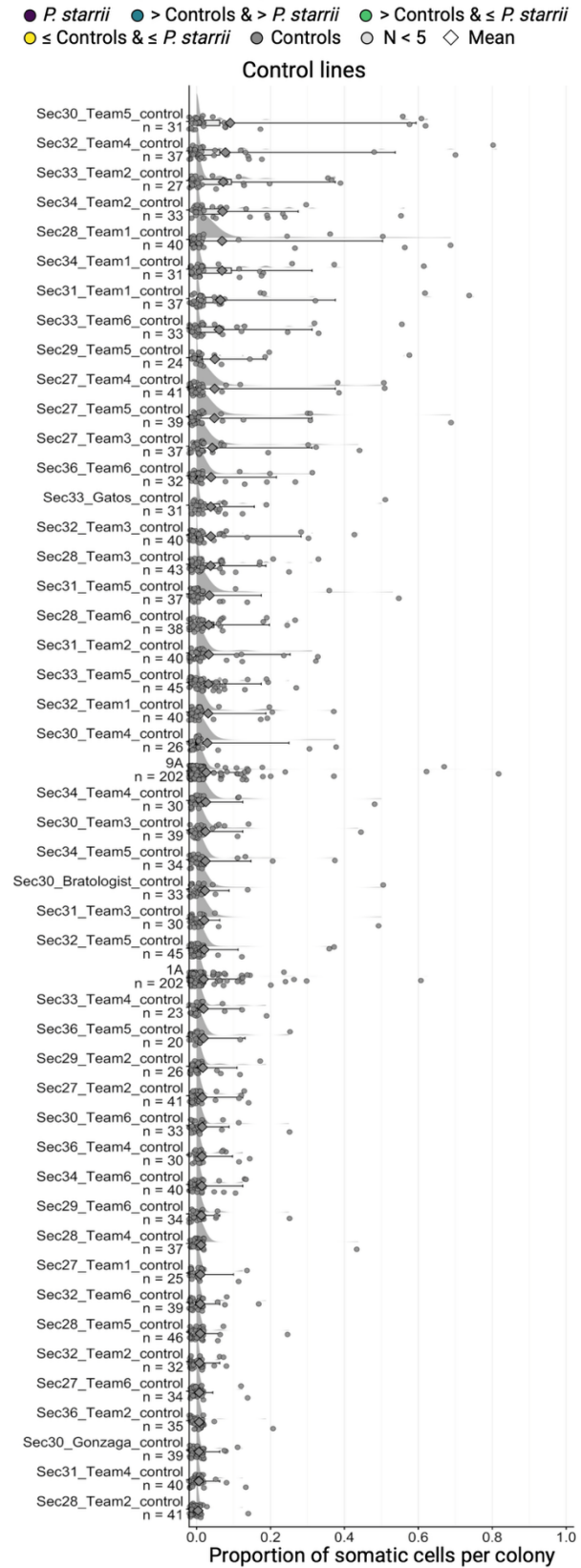

**Figure S9.** The distribution of the proportion of somatic cells per line for colonies of all sizes from all lines. The distribution of somatic cells for *P. starrii* is shown at the top left in dark purple. The mean proportion of somatic cells for *P. starrii* is shown in the dashed purple line. The experimentally evolved lines shown in teal are those in which the mean proportion of somatic cells per colony is significantly greater than that of *P. starrii*. The mean number of somatic cells per colony for each line is shown as a diamond. The lines shown in green are those for which the mean proportion of somatic cells per colony is significantly greater than in pooled controls but not significantly greater than *P. starrii*. Experimentally evolved lines shown in yellow are those for which the proportion of somatic cells per colony is not significantly greater than controls. Controls are shown in gray on the right. Lines with fewer than 5 data points are shown in light gray. *Eudorina* lines are ordered by the mean proportion of somatic cells per colony. Created in https://BioRender.com.

**Table S11.** Summary of morphological changes and comparison to example *Pleodorina* species. Data are drawn from colonies of all sizes (not just 16-celled colonies) and pooled data are combined from the laboratory lines and the CURE. The percent and total proportion of colonies with somatic cells, and the percent of soma per differentiated colony with the standard deviation are shown. Example *Pleodorina* species for which the proportion of somatic cells per differentiated colony overlaps with that of the corresponding lines (or pooled lines) are also shown. The percent soma for each *Pleodorina* species is taken from its species description. Data for *P. japonica* (Nozaki et al., 1989), *P. starrii* (Nozaki et al., 2006), and *P. californica* (Shaw, 1894) are included.

| Treatment or line | Percent (proportion) of colonies with soma | Percent (SD) soma per differentiated colony | Similar *Pleodorina* species | Percent soma per colony in *Pleodorina* |
| --- | --- | --- | --- | --- |
| Pooled controls | 16.9% (340/2012) | 17.5% (0.16) |  |  |
| Pooled lines that didn't evolve soma | 24.8% (224/904) | 19.0% (0.17) | *P. japonica* | 19.5% - 26.6% |
| Pooled lines that evolved more colonies with soma | 51.7% (526/1018) | 25.3% (0.20) | *P. japonica*<br>*P. starrii* | 19.5% - 26.6%<br>25.0% - 37.5% |
| Pooled lines that evolved more soma per colony | 56.0% (432/772) | 30.1% (0.22) | *P. starrii*<br>*P. californica* | 25.0% - 37.5%<br>38% - 50% |
| Section27_Team5 | 91.4% (32/35) | 25.0% (0.15) | *P. japonica*<br>*P. starrii* | 19.5% - 26.6%<br>25.0% - 37.5% |
| 11A5 | 94.1% (64/68) | 40.7% (0.26) | *P. californica* | 38% - 50% |
| Sec33_Gatos | 83.3% (25/30) | 49.3% (0.24) | *P. californica* | 38% - 50% |

#### Genomic changes

##### Mutations

The experimentally evolved line, 11A5, had a significantly higher proportion of non-coding mutations than did the control, 1A. We identified 3,970 fixed mutations unique to 11A5, 5,848 fixed mutations unique to 1A, and 159,455 shared mutations that most likely arose in the years since our culture collection strain has been diverging from the sequenced reference. 11A5 had a significantly smaller proportion of genic mutations than did 1A (Fisher's exact test, odds ratio = 0.43, $p < 0.0001$). 10.4% (414/3970) of 11A5 mutations were genic, while 21.3% (1243/5848) of 1A mutations were genic. 11A5 also had a significantly smaller proportion of CDS mutations than did 1A (11A5: 84/3970; 2.12% vs 1A: 216/5848, 3.69%; Fisher's exact test, odds ratio = 0.56, $p < 0.0001$). However, this was driven by differences in the proportion of intergenic mutations (11A5: 3,556/3,970, 89.57% vs 1A: 4,605/5848, 78.7%), and the two lines did not significantly differ in the proportion of genic mutations in the CDS (11A5: 84/414, 20.29% vs 1A: 216/1243, 17.38%; Fisher's exact test, odds ratio = 1.21, $p > 0.05$). Differences in the total number of mutations and the proportion of intergenic mutations are consistent with the more recent subcloning of 11A5, as 11A5 was subcloned twice after the start of the experiment. Subcloning reduces overall genetic variation and is likely to increase the fixation of mutations via genetic drift.

**Table S12.** Summary of mutations present in only 11A5, only 1A, and shared between the two of them. Total mutations, mutation types, genomic regions, possible CDS consequences, possible splice variants, and structural variants (excluding transposable elements) are summarized. We refer to mutations that are present in only one sample as sample-specific mutations.

| Category | Subcategory | 11A5 | 1A | Shared |
| --- | --- | --- | --- | --- |
| Total mutations |  | 3970 | 5848 | 159455 |
| Mutation type | SNPs | 3483 | 5270 | 155832 |
|  | Indels (true) | 241 | 287 | 226 |
|  | MNVs | 216 | 233 | 114 |
|  | TE insertions | 16 | 47 | 252 |
|  | SVs (non-TE) | 14 | 11 | 3031 |
| Genomic region | Intergenic | 3556 | 4605 | 73401 |
|  | Intronic | 188 | 560 | 41892 |
|  | CDS | 84 | 216 | 17572 |
|  | Promoter | 72 | 280 | 11435 |
|  | 5'UTR | 5 | 15 | 1900 |
|  | 3'UTR | 26 | 49 | 6703 |
|  | Downstream | 28 | 118 | 6521 |
|  | Genic total | 414 | 1243 | 86054 |
|  | % genic | 10.4 | 21.3 | 54.0 |
| **Possible CDS consequences** | Synonymous | 25 | 107 | 9332 |
|  | Missense | 50 | 98 | 7950 |
|  | Nonsense | 1 | 0 | 35 |
|  | Frameshift | 5 | 5 | 6 |
|  | Inframe indel | 1 | 2 | 2 |
|  | Start lost | 0 | 0 | 15 |
|  | Stop lost | 0 | 0 | 19 |
|  | Protein-disrupting total | 6 | 7 | 101 |
| **Possible splice variants** | Canonical (<=2bp) | 0 | 2 | 26 |
|  | Region (3-8bp) | 4 | 9 | 994 |
| **SV types** | DEL | 8 | 6 | 1555 |
|  | INS | 6 | 4 | 1454 |
|  | DUP | 0 | 1 | 6 |
|  | INV | 0 | 0 | 2 |
|  | BND | 0 | 0 | 14 |
|  | TE insertion | 16 | 47 | 252 |

##### Genic changes

**Table S13.** A summary of sample-specific and shared mutations affecting genes in 11A5, 1A. Sample-specific mutations are those that are only present in one line, while shared mutations are mutations that are the same in both lines and may have arisen in culture collection prior to this experiment.

| Category | Subcategory | 11A5 genes | 1A genes | Shared genes |
| --- | --- | --- | --- | --- |
| **Total mutations** |  | 128 | 207 | 79587 |
| **Mutation type** | SNPs | 96 | 179 | 77097 |
|  | Indels (true) | 15 | 11 | 61 |
|  | MNVs | 10 | 6 | 39 |
|  | TE insertions | 3 | 7 | 177 |
|  | SVs (non-TE) | 4 | 4 | 2213 |
| **Genomic region** | Intergenic | 0 | 0 | 0 |
|  | Intronic | 45 | 44 | 38655 |
|  | CDS | 17 | 42 | 16326 |
|  | Promoter | 27 | 75 | 10551 |
|  | 5'UTR | 3 | 5 | 1764 |
|  | 3'UTR | 17 | 8 | 6178 |
|  | Downstream | 9 | 28 | 6082 |
|  | Genic total | 128 | 207 | 79587 |
| **CDS consequences** | Synonymous | 9 | 13 | 8674 |
|  | Missense | 7 | 26 | 7386 |
|  | Nonsense | 0 | 0 | 32 |
|  | Frameshift | 0 | 1 | 5 |
|  | Inframe indel | 0 | 0 | 0 |
|  | Start lost | 0 | 0 | 14 |
|  | Stop lost | 0 | 0 | 17 |

**Table S14.** A summary of genes with sample-specific and shared mutations. If the same gene has different mutations in 11A5 and 1A, it is included in both columns.

| Category | 11A5 | 1A | Shared |
| --- | --- | --- | --- |
| Genes with mutations | 98 | 208 | 3449 |
| Genes with CDS mutations | 19 | 57 | 2607 |
| Genes with intronic mutations | 44 | 93 | 2705 |
| Genes with promoter mutations | 24 | 71 | 2133 |
| Genes with UTR mutations | 13 | 22 | 1509 |
| Genes with missense mutations | 13 | 36 | 1902 |
| Genes with synonymous mutations | 9 | 38 | 2202 |
| Genes with protein-disrupting mutations | 2 | 6 | 91 |
| Genes with splice region variants | 4 | 7 | 706 |
| Genes with TE insertions | 10 | 34 | 180 |
| Genes with TE in CDS | 0 | 3 | 9 |
| Genes with SV (non-TE) | 5 | 7 | 1411 |
| Genes with SV in CDS | 2 | 1 | 179 |

**Table S15.** A summary of sample-specific mutated genes and genes with only mutations that are shared between the two lines. 11A5-specific genes have sample-specific mutations in 11A5 (and lack 1A-specific or shared mutations), 1A-specific genes have sample-specific mutations in 1A (and lack 11A5-specific or shared mutations), and shared-specific genes have identical mutations in 11A5 and 1A (and lack sample-specific mutations).

| Category | 11A5-specific | 1A-specific | Shared-specific |
| --- | --- | --- | --- |
| Genes with mutations | 41 | 57 | 3255 |
| Genes with CDS mutations | 7 | 13 | 2440 |
| Genes with intronic mutations | 14 | 18 | 2546 |
| Genes with promoter mutations | 11 | 19 | 2004 |
| Genes with UTR mutations | 7 | 2 | 1413 |
| Genes with missense mutations | 5 | 8 | 1775 |
| Genes with synonymous mutations | 3 | 7 | 2065 |
| **Genes with protein-disrupting** | 0 | 1 | 84 |
| **Genes with splice region variants** | 1 | 0 | 652 |
| **Genes with TE insertions** | 3 | 7 | 163 |
| **Genes with TE in CDS** | 0 | 1 | 7 |
| **Genes with SV (non-TE)** | 4 | 4 | 1299 |
| **Genes with SV in CDS** | 1 | 1 | 168 |

**Table S16.** 11A5-specific annotated mutated genes. The gene ID, PFAM ID, gene product description, mutations, and possible functional consequences based on mutation locations are provided.

| **Gene ID** | **PFAM** | **Gene product description** | **Mutations** | **Possible functional consequences** |
| --- | --- | --- | --- | --- |
| **EUDS_000603** | Squamosa promoter binding protein (SBP) domain and BTB/POZ domain | SBP proteins are transcription factors involved in developmental regulation; BTB/POZ domains are repressors | CDS SNP and a multi-exon deletion | Disrupted DNA binding domain and SBP loss of function |
| **EUDS_004556** | Thioredoxin-like ferredoxin domain | Electron carrier; may be involved in redox reactions or photosynthesis electron transport | 3'UTR SNP | Regulatory changes |
| **EUDS_005618** | AAA+-type ATPase, SpoVK/Ycf46/Vps4 | ATPase containing an AAA (ATPase Associated with various cellular Activities) domain that may be involved in cell cycle regulation | Downstream SV | Regulatory changes |
| **EUDS_006276** | BRO1/Alix-like; annotation from volvocine homolog | ESCRT system and apoptosis pathways | Intron SV | Alternative splicing changes |
| **EUDS_006857** | Universal stress protein family | May enhance cell survival during stress | Intron TE | Alternative splicing changes |
| **EUDS_009368** | Karyopherin (importin) alpha (SRP1 domain) | Import of proteins into the cell nucleus | 3 promoter SNPs, 1 5'UTR SNP, 8 intron SNPs, 1 downstream SNP, 4 synonymous CDS SNPs | Regulatory and alternative splicing changes |
| **EUDS_011253** | NACHT, WD40, and MLKL_NTD | NACHT and MLKL_NTD domains are involved in apoptosis/programmed cell death | Intron SNP | Alternative splicing changes |
| **EUDS_011729** | K-dependent Na<sup>+</sup>/Ca<sup>+</sup> exchanger | Transport and binding of cations and proteins | 9 promoter SNPs | Regulatory changes |
| **EUDS_014576** | F-box like protein; annotation from volvocine homolog | Protein ubiquitination and degradation | 2 CDS missense indels, 3 3'UTR indels, 1 5'UTR indel, 10 intron indels, 4 downstream indels | Protein-coding, regulatory, and alternative splicing changes |
| **EUDS_014806** | Provisional conjugal transfer protein TrbL; annotation from volvocine homolog | Provisional inner membrane protein that helps form a channel for macromolecule transport | Missense CDS SNP | Protein-coding changes |
| **EUDS_015021** | Sulfite exporter TauE | Transport of anions across the membrane during taurine metabolism; involved | 3'UTR SV | Regulatory changes |
|  |  | in response to oxidative stress |  |  |
| **EUDS_015427** | Serine/threonine protein kinase containing a catalytic domain | Stress-activated kinase that regulates glucose and lipid metabolism and regulates multiple cell processes | Promoter SNP | Regulatory changes |
| **EUDS_016846** | Ion transport protein | Selectively regulates ion transport across the membrane | Intron TE | Alternative splicing changes |
| **EUDS_018116** | Nucleotide-binding domain of heat shock 70 kDa and similar proteins and DnaK domain; annotation from volvocine homolog | Chaperones that assist in protein folding and assembly | 2 intron SNPs and 1 intron SV | Regulatory and alternative splicing changes |
| **EUDS_018198** | SDR family NAD(P)-dependent oxidoreductase | SDR family oxidoreductases are involved in redox sensing and catalyzing numerous activities | Promoter TE | Regulatory changes |
| **EUDS_018202** | Gametolysin peptidase M11; annotation from volvocine homolog | Metalloprotease involved in cell wall degradation | Downstream indel | Regulatory changes |
| **EUDS_019606** | WD40 domain | Coordinates protein-protein interaction and regulates an array of functions | 2 CDS missense SNPs, 5 3'UTR SNPs, 1 5'UTR SNP, 13 intron SNPs, 4 | Protein-coding, regulatory, and alternative splicing changes |
|  |  |  | synonymous<br>CDS SNPs |  |
| **EUDS_020024** | Gametolysin<br>peptidase M11<br>with a zinc-<br>dependant<br>metalloprotease<br>active site and an<br>acidic double-<br>disulfide repeat | Metalloprotease<br>involved in cell wall<br>degradation; may be<br>involved in<br>developmental<br>processes | Downstream<br>indel | Regulatory<br>changes |

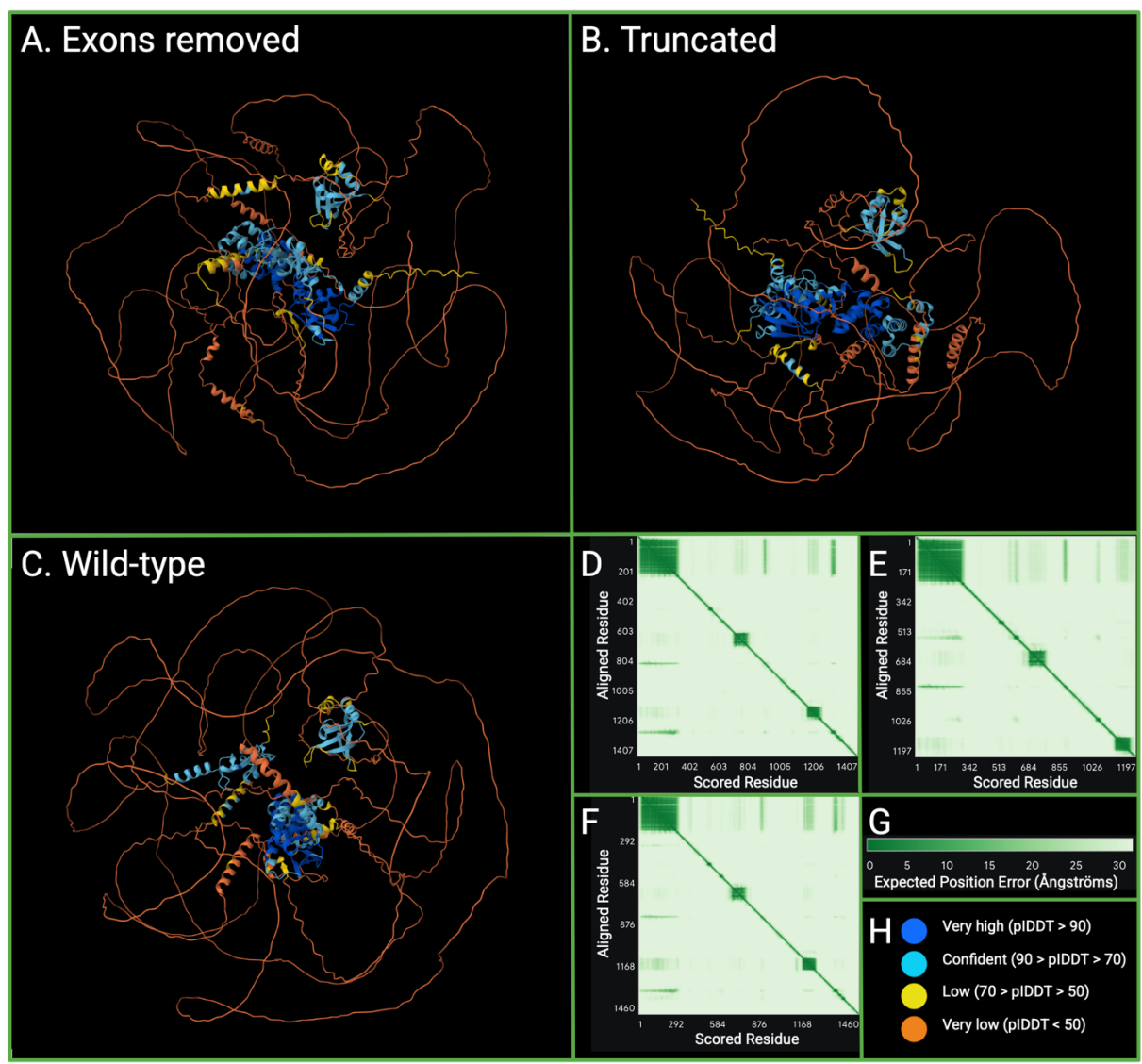

**Figure S10.** Reconstructions of two scenarios resulting from mutations to EUDS\_000603, along with comparisons to wild-type protein structure and predicted alignment errors. Heatmaps show the expected positional error in Ångströms when aligning on residue $y$ to estimate the error at residue $x$. A. EUDS\_000603 with lost exons removed and the rest of the protein translated. Predicted template modeling score = 0.3. B. EUDS\_000603, truncated after the multi-exon deletion. Predicted template modeling score = 0.31. C. Wild-type EUDS\_000603. Predicted template modeling score = 0.3. D. Predicted alignment errors associated with panel A. E. Predicted alignment errors associated with panel B. F. Predicted alignment errors associated with panel C. G. Scale bar for predicted alignment errors (panels D-F). H. Per-atom confidence estimates on a 1-100 scale. Higher values indicate higher confidence. Areas of dark blue in the alignments correspond to very high confidence (pLDDT > 90), light blue corresponds to confident (90 > pLDDT > 70), yellow corresponds to low confidence (70 > pLDDT > 50), and orange corresponds to very low confidence (pLDDT < 50). We generated structure predictions

and predicted alignment errors with AlphaFold 3 (Abramson et al., 2024). Figure created with BioRender.com.

We supplemented our gene annotations by generating 3D protein structures with AlphaFold3 (Abramson et al., 2024) (Figure S10), converting them to PDB files using PyMOL (Yuan et al., 2016), and uploading the PyMOL structures into DALI (Holm, 2020). We also obtained additional annotation information from STRING (Szklarczyk et al., 2019). We used BLAST to identify the *Chlamydomonas* homolog of a translated 11A5-specific mutated gene, EUDS\_000603. We examined the full first-degree STRING network, where the edges indicate both functional and physical protein associations. We required medium-confidence minimum interaction scores. All interactions were supported by known interactions, based on experimentally determined interactions and known databases.

To further investigate the function of gene EUDS\_000603, we examined the predicted functional partners of its *Chlamydomonas reinhardtii* homolog from the STRING database. The translated *Chlamydomonas* homolog of gene EUDS\_000603 likely interacts with two subnetworks that are connected via their interactions with the EUDS\_000603 homolog (identifier A0A2K3E5K7) and with a second protein (identifier A0A2K3DCN5) with 5 ankyrin repeats and a RING domain. GO terms and domain annotations indicate that 4-gene subnetwork is involved in cilium assembly and is part of the microtubule organizing center and cilium while the 5-gene subnetwork is involved in protein ubiquitination and ubiquitin-dependent protein catabolic processes and is part of a cullin-RING ubiquitin ligase complex. Cullin-RING ligases affect many aspects of plant biology, including cell cycle regulation, transcription, and stress responses (Hua & Vierstra, 2011). Full results are available in supplementary material SX. Gene EUDS\_000603, therefore, may be tied to two modules downstream of transcriptional regulation in the response to cold shock (Figure 3): protein degradation and cell cycle regulation.

##### *Comparisons to other volvocine species*

We examined all 15 11A5 and 29 1A sample-specific mutated genes for which we found *Chlamydomonas* homologs. We found that 11A5 had more sample-specific genes (11/15, 73.3% vs 12/29, 41.4%) whose *Chlamydomonas* homologs had cold-induced expression following up to one hour of cold shock, as reported in (Li et al., 2020); this difference was not significant at the $\alpha = 0.05$ level (odds ratio = 3.90, confidence interval = 0.85 – 20.35, $p > 0.05$).

We also compared the proportion of sample-specific mutated genes for which we identified *Volvox* or *Astrephomene* homologs. We found that 11A5 had significantly more sample-specific genes whose homologs had soma-specific expression than did 1A (odds ratio = 3.00, confidence interval = 0.71 – 13.78, $p > 0.05$). 70.6% (12/17) of 11A5 genes had homologs with soma-specific expression, while 44.4% (12/27) of 1A-specific genes had homologs with soma-specific expression. Results are summarized in Table SX8.

**Table S17.** Homolog comparisons for ample-specific mutated genes from experimentally evolved 11A5 vs control 1A. Comparisons conducted with Fisher's exact tests.

| Test | Inclusion criteria | Experimentally evolved 11A5 | Control 1A | Conditional odds ratio | 95% confidence interval | P-value |
| --- | --- | --- | --- | --- | --- | --- |
| Both cold-induced and soma-specific (inclusive) | *Chlamydomonas*, *Volvox*, and *Astrephomene* homologs identified | 10/14 (71.4%) | 7/24 (29.2%) | 5.75 | 1.17-34.50 | < 0.05 |
| Soma-induced or soma-specific (exclusive) | *Chlamydomonas*, *Volvox*, and *Astrephomene* homologs identified | 13/14 (92.9%) | 16/24 (66.7%) | 6.24 | 0.68-309.66 | >0.05 |
| Cold-induced | *Chlamydomonas* homologs identified | 11/15 (73.3%) | 12/29 (41.4%) | 3.77 | 0.85-20.35 | >0.05 |
| Soma-specific | *Volvox* and/or *Astrephomene* homologs identified | 12/17 (70.6%) | 12/27 (44.4%) | 2.92 | 0.71-13.78 | >0.05 |

##### Hypothesized pathway

We propose that the evolution of obligate somatic differentiation in experimentally evolved line 11A5 was predominantly underpinned by regulatory modifications to genes that are part of both cold shock response modules and somatic differentiation modules. This may have facilitated activation of the differentiation module without cold shock to trigger its expression. Additionally, there is strong concordance between annotations of mutated genes and the annotated cold stress response pathway in *Chlamydomonas* and *Arabidopsis*. Moreover, of the 12 mutated genes for which we could identify close *Chlamydomonas* homologs, 9 had *Chlamydomonas* homologs that Li et al. (2020) found were differentially expressed in response to up to an hour of cold shock. We describe the physiological changes we hypothesize occurred in 11A5 based on the mutations we identified.

The initial response to cold shock in *Chlamydomonas* involves the import of calcium across the membrane. In 11A5, the transport of calcium and other ions across the membrane may have been modified, potentially affecting the amount of calcium present in the cell at baseline conditions. We found that there were likely regulatory changes in a voltage gated ion channel (gene 16846) and in a K<sup>+</sup> dependent Na<sup>+</sup>/Ca<sup>+</sup> exchanger (gene 11729). Gene 16846 has a *Volvox carteri* homolog with soma-specific expression and a *Chlamydomonas reinhardtii*

homolog that is upregulated in response to one hour of cold stress. Taken together, this suggests that changes in calcium and other ion transport likely occurred; a modification of what is likely the first step in the response to cold shock that results in the development of plastic somatic cells in wild-type *Eudorina*.

Signal transduction via molecules such as calcium leads to the activation of cold-responsive regulatory factors such as kinases and HSF1. 11A5 has a putative serine-threonine protein kinase (gene 15427) with a promoter SNP. Changes in the expression of this kinase could potentially affect how the downstream cold shock response pathway is activated or modify how sensitive the cold shock response is to the amount of calcium present in the cell. This gene has *C. reinhardtii* homologs that are upregulated in response to an hour of cold shock.

Cold-responsive regulatory factors lead to changes in transcriptional regulation. The transcriptional response to cold shock was likely altered in 11A5. A putative transcription factor with a Squamosa Promoter Binding-Like Protein (SPL) domain and a BTB/POZ domain (gene 603) had a deletion spanning two exons and splice sites, resulting in the removal of the SPL DNA binding domain. SPL domains are found in plant proteins and regulate developmental processes, including the control of flower development and the regulation of vegetative transitions. SPL domain proteins have undergone gene duplication and numerous SPL domain proteins are upregulated in response to cold stress in *C. reinhardtii*. However, the closest *C. reinhardtii* homolog to gene 603 is initially slightly upregulated in response to cold stress and then significantly downregulated after an hour of cold stress. Additionally, *V. carteri* SPL homologs have soma-specific expression. BTB/POZ domains mediate dimerization and can mediate transcriptional repression. Taken together, this indicates that a transcription factor that likely acts as a transcriptional repressor and is downregulated in response to cold shock was lost. This is likely to result in changes in the regulation of the cold shock response, including the potential for causing the constitutive upregulation of downstream components of the cold shock response. The loss of this transcription factor is therefore a key candidate for mediating the transition from environmental to developmental regulation of cellular specialization.

While gene 603 was the only transcription factor mutated in 11A5, it is not the only mutated gene that may regulate the transcriptional response to cold shock. Gene 9368 had 17 predominantly regulatory SNPs and was annotated as SRP1 (karyopherin (importin) $\alpha$), indicating that it is likely involved in regulating intracellular trafficking and secretion. SRP1 facilitates nuclear import signal receptor activity, thereby affecting gene expression. The *C. reinhardtii* homolog is not differentially expressed in response to up to an hour of cold shock but the *V. carteri* homolog has soma-specific expression. Additionally, the *C. reinhardtii* homolog of gene 9368 is localized to the cytoskeleton and is necessary for normal flagellar movement in response to calcium. This is relevant for the regulation of somatic cells because functional flagella are a hallmark of somatic cells, and cell division is regulated by the basal bodies that are flagellar attachment points.

The activation of protective proteins and degrading mechanisms, including proteins regulating redox reactions, typically follows the transcriptional response to cold shock. In 11A5, we observed putative regulatory mutations in 6 genes that we propose fall in this category: genes 6857, 18198, 18116, 4556, 14576, and 15021. There was a TE inserted in the promoter of gene 18198, a putative oxidoreductase. Oxidoreductases regulate redox reactions, and reactive oxygen species (ROS) are important second messengers that mediate the development of sexual differentiation in *Volvox carteri* (Nedelcu, 2009; Nedelcu & Michod, 2006). A *C. reinhardtii* homolog of this gene is upregulated in response to one hour of cold shock, and *V. carteri* homologs exhibit soma-specific gene expression. Gene 4556, with a 3'UTR SNP, is a member of the Thioredoxin-like [2Fe-2S] Ferredoxin family with putative 2Fe-2S cluster binding sites. Proteins with this domain are low-potential electron carriers and homologous domains are found in redox enzymes. A *C. reinhardtii* homolog is upregulated in response to 10 minutes of cold stress. Gene 14576, an F-box protein, had 20 indels; F-box proteins are often involved in protein ubiquitination and degradation. Finally, 15021, annotated as sulfite exporter TauE, had a 3'UTR SV insertion. TauE exports sulfites, thereby contributing to the maintenance of the redox balance.

We also found mutations in two categories of genes that are outside of the canonical cold shock response pathway (Ermilova, 2020). First, we see a set of genes likely involved in cell cycle regulation. 19606, which contains WD40 repeats, has 25 SNPs. WD40 repeats mediate protein-protein interactions, and the *Chlamydomonas* homolog of this protein likely interacts with the flagella and basal bodies and is upregulated in response to cold shock. We found an intron SNP ~50 bp from an exon in gene 11253, which has a WD40 domain as well as a NACHT/MLKL-like domain, indicating that it may be involved in apoptosis and programmed cell death. 6276, which is likely involved in the ESCRT pathway and apoptosis, had a promoter indel. Finally, 5618, a AAA+-type ATPase SpoVK, a likely regulator of cell cycle progression, had a downstream SV.

The final category of genes is cell wall remodeling, one of the functional categories enriched in the expression of soma-specific genes in both *Astrephomene* and *Volvox*. Genes 14806, a provisional GTPase activator, had a missense CDS SNP. We also had two gametolysin peptidase M11 genes with downstream indels: 18202 and 20024. However, since there is a different putative gametolysin peptidase M11 that is mutated in the control, changes in gametolysin peptidase M11 may be a consequence of adaptation to culture collection conditions.

The order in which the genetic changes occurred is unclear. Additionally, it is unknown whether any given sample-specific mutation was involved in adaptation to cold shock, was part of a response to the genetic changes that occurred during repeated cold shock, or was selected for by culture collection conditions. Sample-specific mutations could also be neutral or sequencing artifacts. Regardless, it is apparent that there are modifications to the cold shock response pathway, and that many of the mutated genes are part of both the cold shock response and somatic differentiation expression modules in other volvocine algae species. At least one, and likely more, of these modifications resulted in the stabilization of the plastic response such that somatic cells consistently develop. The most likely candidate shaping the transition from facultative to obligate differentiation is the loss of the DNA-binding domain of gene 603, a transcription factor whose homolog is downregulated in response to cold shock and which may act as a transcriptional repressor. The potential loss of a repressor of the cold shock response may have led to the hypothesized constitutive expression of the cold shock pathway.
